# Nematode-Trapping Fungus *Arthrobotrys oligospora* is Hungry for COQ7 to Alleviate Environmentally Triggered Iron Overload

**DOI:** 10.1101/2024.12.24.630212

**Authors:** Qunfu Wu, Jiao Zhou, Donglou Wang, Songhan Xue, Ling Li, Li Wu, Junxian Yan, Xuemei Niu

## Abstract

Choosing the right and appropriate amount of substances in a changing environment is essential for the growth and survival of all species. We found that elevated oxygen levels significantly enhanced the iron-rich trapping devices of a nematode-trapping fungus (NTF), *Arthrobotrys oligospora*. Interestingly, genomic analysis revealed that all NTFs lacked *coq7*, a key gene involved in ubiquinol (UQ) biosynthesis and iron chelation. Notably, *A. oligospora* utilizes UQ instead of UQ for low-aerobic respiration. Transcriptional, metabolic, mutational, and phenotypic analyses revealed that *A. oligospora* produces a chemotaxonomic class of highly oxygenated arthrobotrins with characteristics similar to UQ skeleton, for high-aerobic respiration. Disruption of arthrobotrin biosynthesis in the Δ*art* mutant not only enhanced UQ biosynthesis, but also triggered trapping device formation and nematicidal activity under elevated oxygen conditions. The addition of UQ to both wild-type and Δ*art* mutants strongly inhibited trapping device development. Remarkably, the iron-rich trapping devices preferentially capture N2 nematodes harboring iron-chelating COQ7 with ferritin properties. Time-calibrated evolutionary analyses, combined with geological data, indicated that the NTF ancestor lost the *coq7* gene after acquiring the arthrobotrin biosynthetic gene cluster during the "superoligotrophy" period, characterized by dramatic shifts in global oxygen levels and temperature changes. Our results demonstrate that oxygen is a critical factor in inducing iron overload, highlighting the origin and role of *coq7* loss in the evolution of predatory behavior of NTFs. Our finding that NTFs capture nematodes for iron chelation therapy, rather than for food, explains a long standing debate about the limited carnivorous ability of trapping devices.

## Introduction

Iron is the most abundant trace metal in organisms and plays essential roles in oxygen transport, the tricarboxylic acid cycle, and electron transport chains [1–4]. However, iron overload can lead to toxicity in all living organisms. Research on iron overload has primarily focused on animal and human models. Excessive iron accumulation can cause tissue damage, particularly in the cardiovascular system. The main causes of iron overload included when iron intake exceeds the body’s ability to regulate it, either through repeated red blood cell transfusions or increased gastrointestinal absorption [5, 6]. Chelation therapy is commonly used to mitigate iron accumulation from blood transfusions by promoting iron excretion through urine and/or feces [7, 8]. A major challenge of chelation therapy is balancing its therapeutic benefits with the risks of over-chelation. Careful dose adjustments are required to avoid excessive chelation of iron as iron levels decrease. To date, few studies have examined the mechanism of iron overload and chelation therapy in other eukaryotes.

A heterogeneous group of organisms within the phylum Ascomycota can differentiate their mycelia into specialized trapping devices to capture nematodes as a primary nutritional source under nutrient-deficient conditions [9–11]. These fungi, classified as nematode-trapping fungi (NTFs), form a monophyletic group within the class Orbiliomycetes and develop various trapping structures, including adhesive networks, adhesive knobs, and constricting rings [11]. Trapping devices have long been regarded as a hallmark of the transition of NTFs from a saprophytic to a predatory lifestyle [12]. Nematodes, the most abundant animals on Earth, occupy all trophic levels within the soil food web [13]. Among them, plant-pathogenic nematodes are highly destructive pests, infecting over 4,000 plant species and causing global crop losses exceeding $100 billion annually [14]. NTFs are recognized as effective natural antagonists that help regulate nematode populations across various ecosystems [15]. However, these carnivorous fungi are neither abundant nor widely distributed in many environments where small animal prey are readily available [9]. It has been hypothesized that the evolution of NTFs is characterized by a degeneration of carnivorous abilities [16]. The mechanisms underlying the limited carnivorous ability of NTF trapping devices remain poorly understood.

Recent studies have shown that the absence of desferriferrichromes or the addition of iron to the predominant nematode-trapping fungus (NTF) *Arthrobotrys oligospora* can induce the spontaneous formation of trapping devices in the absence of nematode prey [17]. Biochemical and chemical analyses have demonstrated that the trapping devices of *A. oligospora* contain more iron than the mycelia, as evidenced by the iron-rich, electron-dense bodies accumulated in the trapping devices [17, 18]. Genomic, evolutionary, and functional analyses further indicate that NTFs have evolved specialized trapping devices to sequester excess iron, compensating for the lack of the Ccc1-mediated vacuolar iron detoxification mechanism, which is conserved in most fungi [18]. Surprisingly, ambient temperature has been identified as a key factor driving the evolution of iron storage strategies and the origin of trapping devices, as iron content and composition in fungal mycelia are inversely correlated with temperature [18]. However, it remains unclear whether other factors contribute to the formation of trapping devices and how the iron-rich nature of these structures relates to the carnivorous ability of NTFs.

Interestingly, *A. oligospora* produces a class of chemotaxonomic meroterpenoids, including arthrobotrisins, which feature a sesquiterpenyl unit linked to an epoxy-cyclohexanol ring [19]. All arthrobotrisin metabolites are derived from a key hybrid precursor, farnesyl toluquinol, which shares a distinctive feature with ubiquinol-3 (UQ_3_): the farnesyl chain is attached to a benzoquinol structure [20]. Ubiquinols (UQs) are essential electron carriers in aerobic respiration, serving as redox intermediates in both aerobic bacteria and eukaryotes [21, 22]. UQs are classified into three subfamilies based on their prenyl chain length: short-chain (C15–C25 for UQ_3_–UQ_5_), medium-chain (C30–C35 for UQ_6_–UQ_7_), and long-chain (C40–C50 for UQ_8_–UQ_10_) [23]. Notably, a series of oxidation processes are critical for converting farnesyl toluquinol into arthrobotrisins, as each arthrobotrisin incorporates at least four oxygen atoms [19, 20, 24–28]. The absence of genes involved in these oxidation processes—specifically the dehydrogenase gene *Ao274* and the *P450* genes *Ao278*, *Ao280*, and *Ao282* in the *art* gene cluster—significantly enhances trapping device formation in the mutants [20, 26–28]. However, the underlying mechanism linking the inverse relationship between arthrobotrisin biosynthesis and trapping device formation via oxygenation remains unexplored.

Oxygen, along with iron, is one of the most abundant elements and plays a crucial role in biological processes. Specifically, the red color of blood cells is the result of the interaction between oxygen and iron. All eukaryotic organisms must maintain oxygen homeostasis, as oxygen is essential for aerobic respiration but also poses significant risks to cellular function [29, 30]. Traditionally, it has been assumed that biological defenses against hyperoxia are less robust than those against hypoxia, since eukaryotes may not have encountered supraphysiological oxygen levels during evolution [29, 30]. However, geological evidence suggests that rising atmospheric oxygen levels were a key driver in the origin and evolution of aerobic life [31–34]. Global oxygen levels gradually increased from around 1.78% in the early Ordovician (approximately 477 million years ago (Mya)) to about 19.88% in the middle Silurian (around 427 Mya) [35]. At various points in history, atmospheric oxygen levels surpassed the current 21%, reaching about 30% during the Carboniferous (around 300 Mya) and Cretaceous (around 100 Mya) periods [35]. To date, limited research has been conducted on the role of oxygen in the evolution of metabolism, genomes, and morphology in NTFs.

As an initial investigation, we examined the role of oxygen in the formation of iron-rich trapping devices in *A. oligospora*. Notably, reduced oxygen levels resulted in a decrease in the formation of trapping devices. Genomic analysis unexpectedly revealed that only 25 out of 2057 fungi, including all NTFs, had lost the *coq7* gene, which encodes an iron-dependent hydroxylase involved in UQ biosynthesis [36]. The COQ7 protein is also a membrane-bound di-iron protein belonging to a unique subgroup of the ferritin family, involved in iron chelation [37–39]. Metabolic analysis of *A. oligospora* showed that the fungus utilizes the atypical ubiquinone UQ_8_ for low-aerobic respiration, in contrast to the more common UQ_9_ and UQ_10_ found in most fungi. Supplementation of *A. oligospora* with either UQ_10_ or the exogenous *coq7* gene from yeast strongly inhibited the development of trapping devices. Remarkably, we observed that the trapping devices of *A. oligospora* were effective in killing N2 nematode strains containing COQ7, but not those lacking it, suggesting that the iron-chelating COQ7 in nematodes is the primary target of the fungus. By correlating oxygen content across the Phanerozoic eon with the evolutionary trajectory of the *art* gene cluster in NTFs and non-NTFs, we found that the dynamics of the *art* gene cluster closely mirror fluctuations in oxygen levels. Physiological and phenotypic analyses of the fungal WT and mutant strains deficient in arthrobotrisin biosynthesis demonstrated that arthrobotrisin production is essential for fungal adaptation to oxygen-rich environments and inhibits the formation of oxygen-induced trapping devices in *A. oligospora*. These results indicate that oxygen is a critical factor in the metabolic evolution of NTFs, influencing both the formation of trapping devices and their nematicidal activity. Our findings suggest that NTFs utilize chelation therapy to mitigate iron overload and that their limited nematicidal activity is necessary to avoid excessive chelation.

## Results

### 1. Oxygen is a factor in inducing trapping devices in *A. oligospora*

Two oxygen conditions were established to assess the role of oxygen in fungal growth, trapping device formation, and nematicidal activity: 21% atmospheric oxygen (normal) and approximately 10% atmospheric oxygen (low). Under low oxygen conditions, the fungal strains exhibited smaller aerial mycelial colonies and developed significantly larger hollow areas compared to those grown under normal oxygen levels (Figs. 1A–1B), indicating that reduced oxygen levels inhibited aerial mycelial growth. After nematode treatment, the fungal strain grown under normal oxygen conditions produced trapping devices, whereas those grown under low oxygen conditions did not (Figs. 1A and 1C). Additionally, the WT strain under normal oxygen conditions exhibited strong nematode-trapping activity, while strains grown under low oxygen conditions did not (Fig. 1D). To rule out the possibility that these differences were due to unequal growth of aerial mycelia under the two oxygen conditions, we selected strains with similar mycelial growth under both conditions for further analysis of trapping device formation. Specifically, the aerial mycelia grown under normal oxygen were one day younger than those grown under low oxygen. Previous studies have suggested that mycelial senescence or nutrient deficiency can easily trigger the formation of trapping devices [17, 40]. However, we observed that the younger aerial mycelia under normal oxygen conditions developed trapping devices, while those under low oxygen conditions did not.

**Fig 1.**
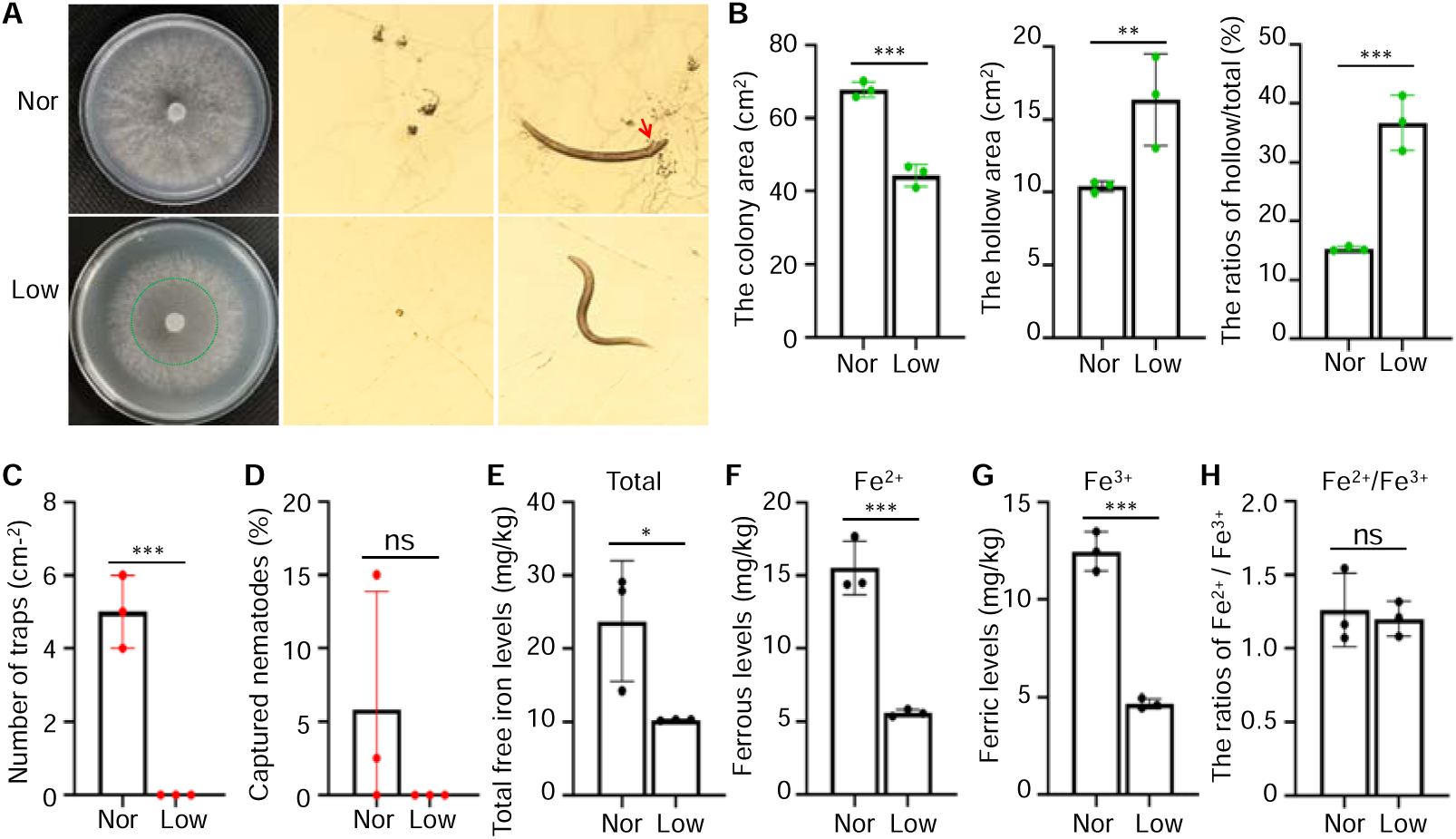
Effects of oxygen on the formation of trapping devices and nematicidal activity in *A. oligospora*. (**A**) Phenotypic analysis of wild-type (WT) under normal (atmospheric oxygen, approximately 21%) and low oxygen (approximately 10%) conditions was performed, including colony growth, trapping device formation, and nematicidal activity. Nor: normal oxygen condition. Low: low oxygen condition. Red arrow: nematode is captured by trapping device. (**B**) Comparison of aerial mycelial areas of WT under normal and low oxygen conditions. (**C**) Comparison of trapping devices of WT induced by nematodes under normal and low oxygen conditions. (**D**) Comparison of nematicidal activity of WT between normal and low oxygen conditions. (**E**−**G**) Comparison of total free iron (E), ferrous (F), and ferric (G) contents in WT under normal and low oxygen conditions. (**H**) Comparison of the ferrous/ferric ratios in WT under normal and low oxygen conditions. Significance was tested using an unpaired t test (\**P* < 0.05, \*\**P* < 0.01, \*\*\**P* < 0.001, ns = not significant).

In a previous study, we demonstrated that *A. oligospora* mycelia develop trapping devices to store excess iron [18]. To evaluate the role of oxygen in regulating iron levels in fungal mycelia, we measured the total free iron content under two oxygen conditions. As anticipated, mycelia grown under normal oxygen conditions contained significantly higher levels of total free iron, including both ferrous and ferric ions, compared to those grown under low oxygen conditions (Figs. 1E–1G). The levels of ferrous, ferric, and total free iron under normal oxygen were more than double those under low oxygen, reflecting the difference in oxygen levels between the two conditions. However, the ratios of Fe(II)/Fe(III) in the mycelia were similar under both conditions (Fig. 1H), consistent with the need to maintain the redox potential of the Fe(II)/Fe(III) ratio in organisms [41]. These results suggest that higher oxygen levels significantly promote the formation of trapping devices by increasing the free iron content in the mycelia.

## 2. All the NTFs lack the *coq7* gene

Previous studies have shown that most fungi utilize UQs with long prenyl chain lengths between 9 and 10 for aerobic respiration [21, 42, 43], and the synthesis of UQ_9_ and UQ_10_ requires at least thirteen proteins, including COQ1–COQ11, YAH1, and ARH1 [43, 44]. Among these, COQ7 has dual functions, participating in both UQ biosynthesis and iron chelation as ferritin [37–39]. To investigate the distribution of these proteins across the fungal kingdom, we used protein sequences of COQ1–COQ11, YAH1, and ARH1 from *Saccharomyces cerevisiae* as queries. These sequences were employed to search for orthologs using BLASTp against a custom database containing predicted protein sequences from 2,057 high-quality fungal genomes. Analysis of the presence and absence of UQ biosynthesis genes revealed that, except for *coq9* (56.73%, 1,167/2,057) and *coq10* (38.89%, 800/2,057), all other genes, including *coq1* (99.85%, 2,054/2,057), *coq2* (99.71%, 2,051/2,057), *coq3* (99.71%, 2,051/2,057), *coq4* (99.76%, 2,052/2,057), *coq5* (99.81%, 2,053/2,057), *coq6* (99.81%, 2,053/2,057), *coq7* (98.78%, 2,032/2,057), *coq8* (99.85%, 2,054/2,057), *coq11* (98.06%, 2,017/2,057), *arh1* (99.90%, 2,055/2,057), and *yah1* (99.56%, 2,048/2,057), are highly conserved across most fungal genomes, with presence frequencies exceeding 98.06% (Fig. 2A and Table S1). This suggests that these eleven genes are essential for UQ biosynthesis and for maintaining metabolic efficiency and viability in fungi. Interestingly, only 25 of 2,057 fungal genomes were missing the critical *coq7* gene, including 9 from Ascomycota, 12 from Basidiomycota, 2 from Neocallimastigomycota, and 2 from Zoopagomycota. Notably, all 9 Ascomycota fungi were NTFs.

**Fig 2.**
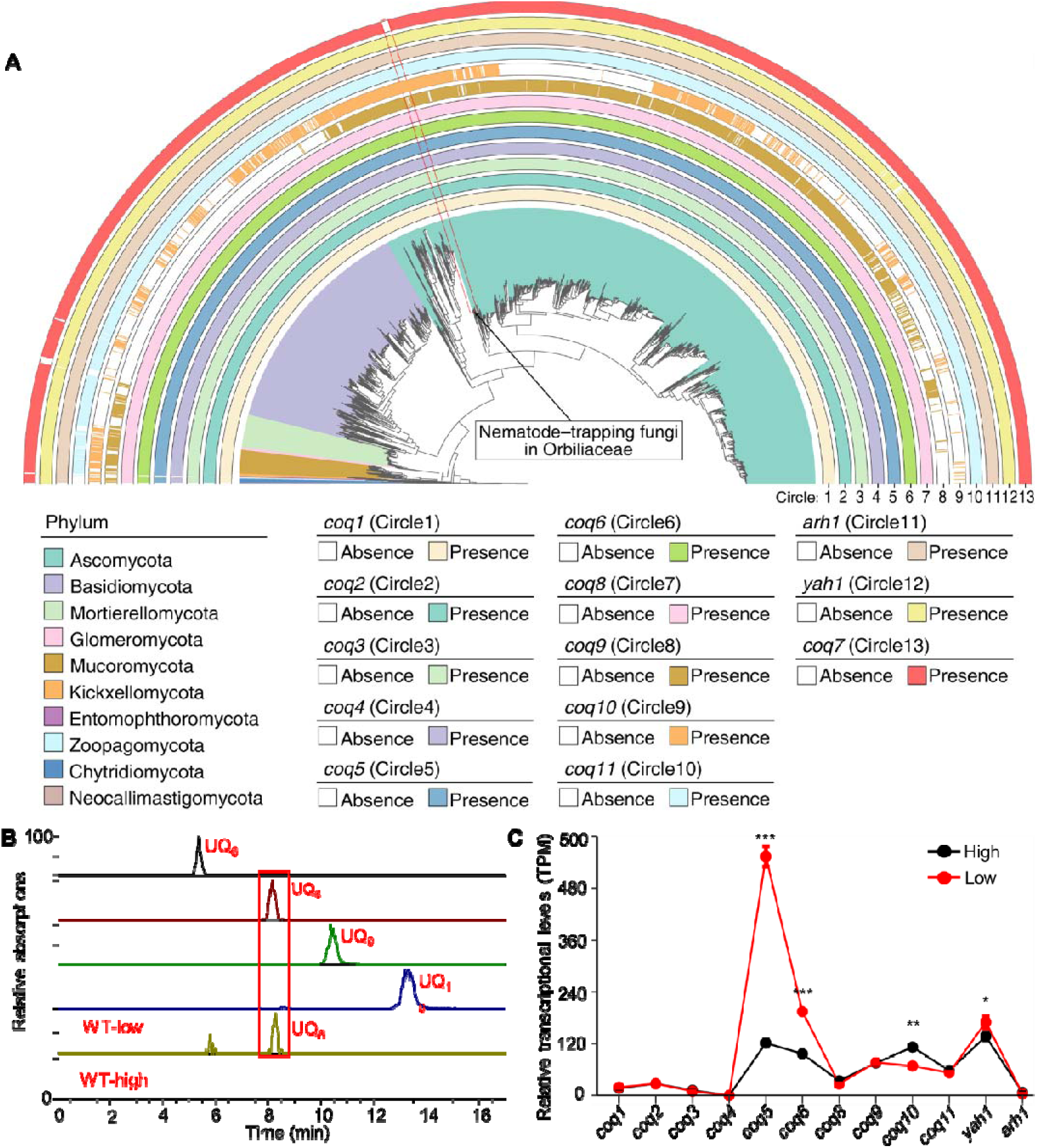
NTFs lack the iron chelating COQ7 for the biosynthesis of UQ_9_ and UQ_10_ for low aerobic condition. (**A**) Distribution patterns of UQ synthesis-related proteins across the fungal kingdom. The phylogenetic tree was constructed using 750 universal single-copy marker genes extracted from 2057 high-quality fungal genomes downloaded from the JGI MycoCosm database. The fungal groups analyzed including Ascomycota (1361 genomes), Basidiomycota (540 genomes), Mucoromycota (57 genomes), Glomeromycota (7 genomes), Mortierellomycota (67 genomes), Kickxellomycota (6 genomes), Chytridiomycota (11 genomes), Neocallimastigomycota (2 genomes), Entomophthoromycota (4 genomes), and Zoopagomycota (2 genomes). The presence or absence of the 13 UQ biosynthesis-related proteins (COQ1–COQ11, YAH1, and ARH1) across various fungal phyla were determined by searching for orthologs of those protein against a custom database containing all predicted protein sequences from 2057 high-quality fungal genomes via BLASTP. The presence or absence of the 13 proteins were mapped onto the tree of fungi kingdom to visualize the distribution patterns of each protein. Each circle (1−13) outside the tree represents the distribution patterns of one of the 13 proteins. Each white block within a circle represents the absence of the corresponding protein in the respective genome and another color represents the presence. Genomes of nematode–trapping fungi in Orbiliaceae are marked in a red box. (**B**) Metabolic analysis of the UQs production in *A. oligospora* under high and low aerobic conditions. (**C**) Transcriptional analysis of 12 UQs biosynthesis-related genes in *A. oligospora* under high and low aerobic conditions. Significance was tested using wald test (\**P* < 0.05, \*\**P* < 0.01, \*\*\**P* < 0.001).

Metabolic analysis of *A. oligospora* under two aerobic conditions revealed the absence of both UQ_9_ and UQ_10_ in the metabolic profiles of the WT strain, consistent with the lack of the *coq7* gene in *A. oligospora* (Fig. 2B). Instead, UQ_8_ was present in the fungus cultured under low aerobic conditions, while its levels were barely detectable under high aerobic conditions (Fig. 2B). Notably, transcriptional analysis revealed no significant changes in the expression of 8 UQ biosynthetic genes between high and low aerobic conditions. In contrast, *coq5, coq6* and *yah1* were significantly down-regulated under high aerobic conditions compared to low aerobic conditions (Fig. 2C). These results suggest that *A. oligospora* utilizes UQ_8_, rather than the more common UQ_9_ and UQ_10_, for respiration under low aerobic conditions.

### 3. Trapping devices of *A. oligospora* prefer *coq7*-containing nematodes

We hypothesized that *A. oligospora* might require UQs with long prenyl chains (9–10 units) under normal oxygen conditions. To test this, we supplemented *A. oligospora* with UQ_10_ at concentrations of 10 and 50 μM under normal oxygen conditions. These concentrations have been shown to have no negative effects on human cells [45], with solvent-only serving as the control. Additionally, the absence of *coq7* in *A. oligospora* led us to hypothesize that the fungus may rely on trapping devices to prey on nematodes as a means of acquiring COQ7. The *clk-1* gene in the nematode *Caenorhabditis elegans* encodes a 187-residue polypeptide homologous to yeast *coq7*, which is essential for UQ biosynthesis in yeast [46, 47]. Therefore, we used the nematode mutant Δ*clk-1* and its WT strain N2 to evaluate trapping device formation and nematicidal activity in *A. oligospora*. Notably, the Δ*clk-1* mutant showed a similar effect on trapping device induction as the N2 nematodes, even slightly higher when *A. oligospora* was treated with 0 and 10 μM of UQ_10_ (Figs. 3A–3B), suggesting that the genetic defect in the nematodes did not affect trapping device formation in the fungus. Meanwhile, UQ_10_ at a concentration of 50 μM strongly inhibited the formation of trapping devices in *A. oligospora*, but had no effect at 10 μM (Figs. 3A–3B).

**Fig 3.**
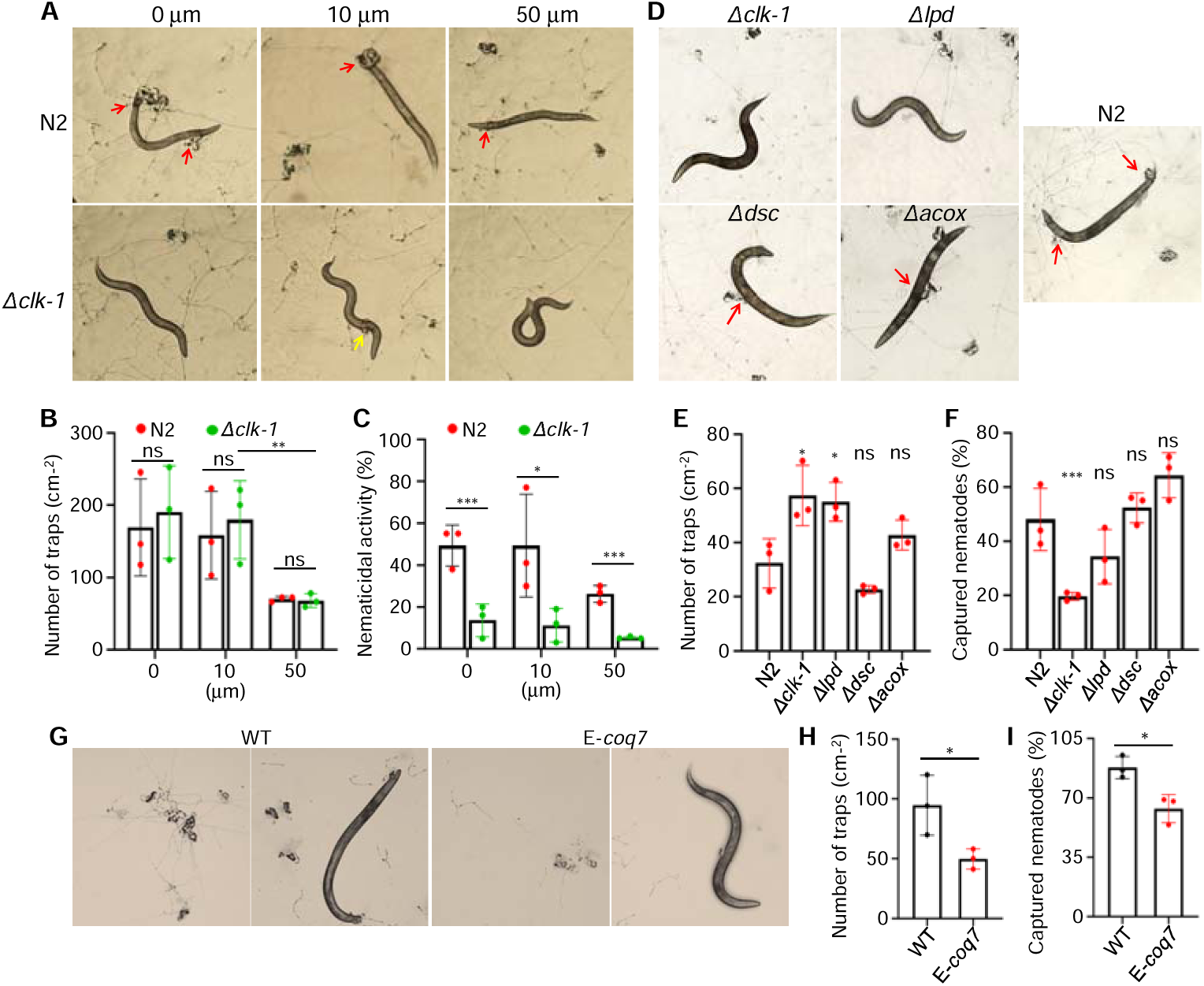
Evaluation of the effects of UQ_10_ and COQ7 on the formation of trapping devices and their nematicidal ability in *A. oligospora*. (**A**) Pictures of the formation of trapping devices in *A. oligospora* treated with two different nematodes WT N2 and mutant Δ*clk-1* in the presence of UQ_10_ at concentrations of 0, 10 and 50μm. 0: with solvent; 10μm: with 10μm UQ_10_; 50μm: with 50μm UQ_10_. Red arrow: trapping device is capturing a nematode. Yellow arrow: nematode is escaping from trapping device. (**B**) Comparison of trapping devices in *A. oligospora* induced by two different nematodes WT N2 and mutant Δ*clk-1* in the presence of UQ_10_, revealed that UQ_10_ at a concentration of 50μm strongly inhibited the formation of trapping devices (**C**) Comparison of nematicidal ability of the trapping devices in *A. oligospora* induced by two nematodes WT N2 and the mutant Δ*clk-1* in the presence of UQ_10_ indicated that the trapping devices preferred to capture the nematodes N2 over the mutant Δ*clk-1*. (**D**) Pictures of the formation of trapping devices in *A. oligospora* induced by five different nematodes, including nematode N2, and the nematode mutants Δ*clk-1*, Δ*lpd*, Δ*dsc*, and Δ*acox*. (**E**) Comparison of trapping devices in *A. oligospora* induced by five different nematodes revealed that the nematode mutants Δ*clk-1* and Δ*lpd* induced the most trapping devices in *A. oligospora*. (**F**) Comparison of nematicidal ability of trapping devices in *A. oligospora* induced by five different nematodes indicated that the nematode mutant Δ*clk-1* is the last prey the trapping devices prefer to capture. (**G**−**I**) Comparison of trapping devices and nematicidal activities between WT and the mutant E-*coq7* with harboring an exogenous *coq7* gene from yeast. Significance was tested using an unpaired t test (\**P* < 0.05, \*\**P* < 0.01, \*\*\**P* < 0.001, ns = not significant).

Notably, we observed that the trapping devices of *A. oligospora* captured significantly more N2 nematodes than the Δ*clk-1* mutant nematodes (Fig. 3C), despite the Δ*clk-1* mutant exhibiting a general slowdown in behavioral rates compared to N2 nematodes [46, 48, 49]. In fact, most Δ*clk-1* nematodes were able to escape from the trapping devices and displayed a vibrant, curved shape, in contrast to the stiff, captured N2 nematodes (Fig. 3A). The ability of *A. oligospora* to capture N2 nematodes was nearly three times greater than that of the Δ*clk-1* nematodes (Fig. 3C). To further investigate the fungal trapping device’s preference for COQ7, we selected three additional nematode mutants deficient in genes encoding Acyl-CoA oxidase (acox), paired-like homeobox transcription factor/defecation suppressor (dsc), and bridge-like lipid transfer protein (lpd), respectively. Among these, the genes *acox* and *lpd* are also absent in all NTFs. The Δ*acox* and Δ*lpd* nematode mutants exhibited locomotor sluggishness comparable to that of the Δ*clk-1* mutant. Phenotypic analysis indicated that the Δ*clk-1* and Δ*lpd* mutants had the strongest effects on inducing the formation of trapping devices, while the Δ*dsc* mutant had the least (Figs. 3D–3E). However, the Δ*dsc* nematode mutant was the second-most-preferred prey for the fungus after the Δ*acox* nematode mutant (Fig. 3F). The trapping devices’ ability to kill the Δ*acox* nematode mutant was twice as high as that for the Δ*clk-1* nematode mutant (Fig. 3F). These results suggest that the trapping devices had the weakest nematode-capturing ability toward the Δ*clk-1* nematode mutant.

To verify the role of COQ7 in the formation of trapping devices and nematicidal activity, the *coq7* gene (819 bp) was directly cloned from *Saccharomyces cerevisiae* BJ5464 and inserted into the *A. oligospora* WT strain to generate the E*-coq7* mutant (Figs. S1A–S1B). Transcriptional analysis confirmed that the *coq7* gene was successfully expressed in the E*-coq7* mutant (Fig. S1C). Metabolic analysis revealed that only UQ_8_, and not UQ_9_ or UQ_10_, was present in the E-*coq7* mutant (Fig. S2), suggesting that the exogenous *coq7* did not participate in UQ biosynthesis in *A. oligospora*. Phenotypic analysis showed that the E-*coq7* mutant exhibited significantly reduced trapping device formation and nematicidal activity compared to the WT strain (Figs. 3G–3I). Given that trapping devices are rich in iron and COQ7 in nematodes functions as a ferritin, the limited nematicidal activity observed in the trapping devices of the fungus appears to align with the chelation therapy in human. Namely, the fungus must carefully control nematode doses to prevent a decrease in iron levels due to excessive chelation.

### 4. *Art* gene cluster is required for elevated aerobic conditions

Due to the lack of coq7 and the low transcriptional levels of UQ biosynthesis under relatively high aerobic conditions, we wondered whether NTFs employ an unknown strategy for normal aerobic respiration. Previous research has shown that deficiency in the highly oxygenated arthrobotrisin resulted in a significant increase in trapping device formation in *A. oligospora* [40]. To investigate the role of the arthrobotrisins in fungal aerobic respiration, the *A. oligospora* WT strain was cultivated under shaking and static conditions. The primary difference between these conditions lies in oxygen transfer and mixing: shaking enhances oxygen availability by increasing the surface area between the liquid culture and the air, facilitating gas exchange and promoting uniform nutrient mixing, which leads to better aeration and more efficient microbial growth, particularly for aerobic organisms [50–52]. Transcriptional analysis revealed that biosynthetic genes in the *art* gene cluster, including *Ao283*–*Ao276* and *Ao283*–*Ao274*–Ao273, were significantly up-regulated under shaking conditions compared to static conditions, while the adjacent non-biosynthetic genes *Ao275* and *Ao284*–*Ao286* remained unchanged (Fig. 4A). Further metabolic analysis demonstrated that arthrobotrisins significantly accumulated in the fungus grown under shaking conditions but were almost absent under static conditions (Fig. 4B).

**Fig 4.**
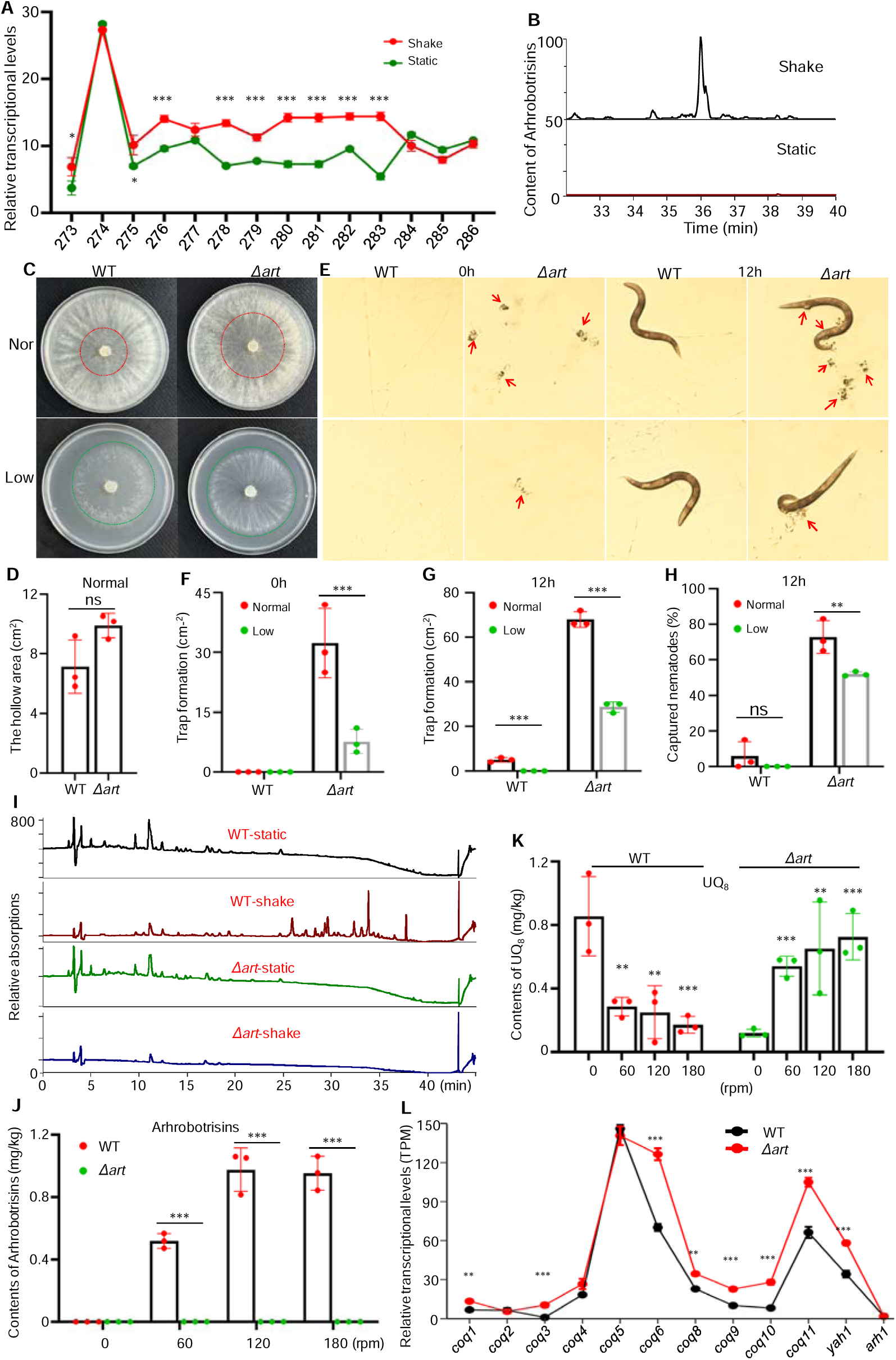
Evaluation of the role of arthrobotrisins in aerial growth, trapping devices, and nematicidal ability in *A. oligospora* and fungal metabolic response to elevated oxygen. (**A**) Transcriptional levels of the genes in the *art* gene cluster in *A. oligospora* WT under shaking and static conditions. (**B**) Metabolic analysis of the major arthrobotrisins in *A. oligospora* under shaking and static conditions. (**C**) Colony growths of *A. oligospora* WT and the mutant Δ*art* under normal and low oxygen conditions. (**D**) Comparison of the hollow area in aerial growth between WT and the mutant Δ*art* under normal oxygen condition. (**E**) Pictures of the formation of trapping devices and nematicidal activities of WT and the mutant Δ*art* under normal and low oxygen conditions within 12h. (**F**) Comparison of trapping devices between WT and the mutant Δ*art* under normal and low oxygen conditions within 0h after nematode treatment. (**G**) Comparison of trapping devices between WT and the mutant Δ*art* under normal and low oxygen conditions within 12h after nematode treatment. (**H**) Comparison of nematicidal activities between WT and the mutant Δ*art* under normal and low oxygen conditions within 12h after nematode treatment. (**I**) Metabolic analysis of *A. oligospora* WT and the mutant Δ*art* under shaking and static conditions. (**J**) Comparison of the contents of the major arthrobotrisins in WT and the mutant Δ*art* under four different shaking conditions of 0 rpm−180rpm. (**K**) Comparison of the contents of UQ_8_ in WT and the mutant Δ*art* under four different shaking conditions of 0 rpm−180rpm. (**L**) Transcriptional levels of the genes responsible for the UQ biosynthesis in *A. oligospora* in the mutant Δ*art* under shaking and static conditions. Significance was tested using an unpaired t test (\**P* < 0.05, \*\**P* < 0.01, \*\*\**P* < 0.001, ns = not significant).

To analyze the impact of arthrobotrisins on fungal morphology and phenotype, we constructed a mutant strain, Δ*art*, lacking arthrobotrisin biosynthesis, based on a previous study [40]. The Δ*art* mutant and WT strains were cultivated on plates under normal (21% atmospheric oxygen) and low oxygen conditions (approximately 10% atmospheric oxygen). Notably, the Δ*art* mutant showed significantly larger hollow areas in the colony center compared to the WT strain under normal oxygen conditions (Figs. 4C–4D), while both strains exhibited similar colony growth under low oxygen levels. This suggests that the absence of arthrobotrisins leads to reduced aerial mycelial formation under normal oxygen conditions. Interestingly, the Δ*art* mutant formed trapping devices without nematode treatment under both aerobic conditions (Fig. 4E), while WT not. The Δ*art* mutant produced significantly more trapping devices under normal aerobic conditions compared to low oxygen conditions (Fig. 4F). Similar results were observed for the Δ*art* mutant when treated with nematodes under both aerobic conditions. After 12 hours of nematode treatment, both the Δ*art* mutant and WT strains developed significantly more trapping devices under normal aerobic conditions compared to low oxygen conditions (Figs. 4E and 4G). Furthermore, the Δ*art* mutant exhibited a greater increase in trapping device formation and nematicidal activity under both aerobic conditions compared to the WT strain (Fig. 4H).

We compared the metabolic profiles of WT and Δ*art* mutant strains under shaking and static conditions. Principal component analysis (PCA) revealed that, under static conditions, the WT and Δ*art* mutant strains clustered together, while they did not cluster together under shaking conditions (Fig. S3). Remarkably, the WT strain exhibited markedly different metabolic profiles under shaking versus static conditions, whereas the Δ*art* mutant did not (Fig. 4I). Additionally, the metabolic profiles of the Δ*art* mutant under both shaking and static conditions were quite similar to those of the WT strain under static conditions, suggesting that arthrobotrisins play a crucial role in the fungal metabolic response to elevated oxygen conditions.

To assess whether the contents of the major arthrobotrisins were influenced by oxygen levels, we cultured the WT and Δ*art* mutant strains under four different shaking conditions: 0, 60, 120, and 180 rpm. Under shaking conditions at 180 rpm, the dissolved oxygen levels are much higher due to enhanced mixing, which increases the oxygen transfer rate from the headspace to the liquid medium [53, 54]. Metabolic analysis showed that the content of the major arthrobotrisins was highest in the WT strain under shaking speeds of 120 and 180 rpm (Fig. 4J). In the shaking speed range of 0–120 rpm, the content of the major arthrobotrisins increased with the shaking speed (Fig. 4J), indicating that their levels are significantly oxygen-dependent. Meanwhile, we observed that UQ_8_ content in the WT strain decreased with increasing shaking speed from 0 to 180 rpm (Fig. 4K). However, in the Δ*art* mutant, UQ_8_ content increased with increasing shaking speed over the same range (Fig. 4K). Transcriptional analysis revealed that UQ biosynthetic genes were significantly up-regulated in the Δ*art* mutant compared to the WT strain (Fig. 4L), indicating that the absence of arthrobotrisins may trigger the production of UQ_8_ in response to elevated oxygen levels.

### 5. The mutant **Δ***art* is hungrier for UQ_10_ and COQ7

Since *A. oligospora* uses arthrobotrisins under elevated oxygen conditions, and the Δ*art* mutant, lacking arthrobotrisins, formed significantly more trapping devices and exhibited stronger nematicidal activity than the WT, we hypothesized that the Δ*art* mutant may have a higher demand for UQ_10_ and COQ7 compared to the WT strain. To test this hypothesis, we supplemented the Δ*art* mutant with UQ_10_ at concentrations of 10, 50, and 500 μM under normal oxygen conditions. Two controls were used: a solvent-only control and an untreated control. Since the Δ*art* mutant can spontaneously develop some trapping devices, we induced the formation of trapping devices both in the presence and absence of nematodes. In both cases, the formation of trapping devices in the Δ*art* mutant treated with all concentrations of UQ10 was significantly reduced compared to the controls (Figs. 5A–5C), indicating that the inhibitory effect of UQ_10_ on trapping device formation was concentration-dependent. Notably, at a concentration of 10 μM UQ_10_, the Δ*art* mutant showed slightly reduced trapping device formation (Figs. 5A–5C), whereas the WT strain did not (Fig. 3B). This suggests that the Δ*art* mutant has a higher demand for UQ_10_ than the WT strain in controlling trapping device formation.

**Fig 5.**
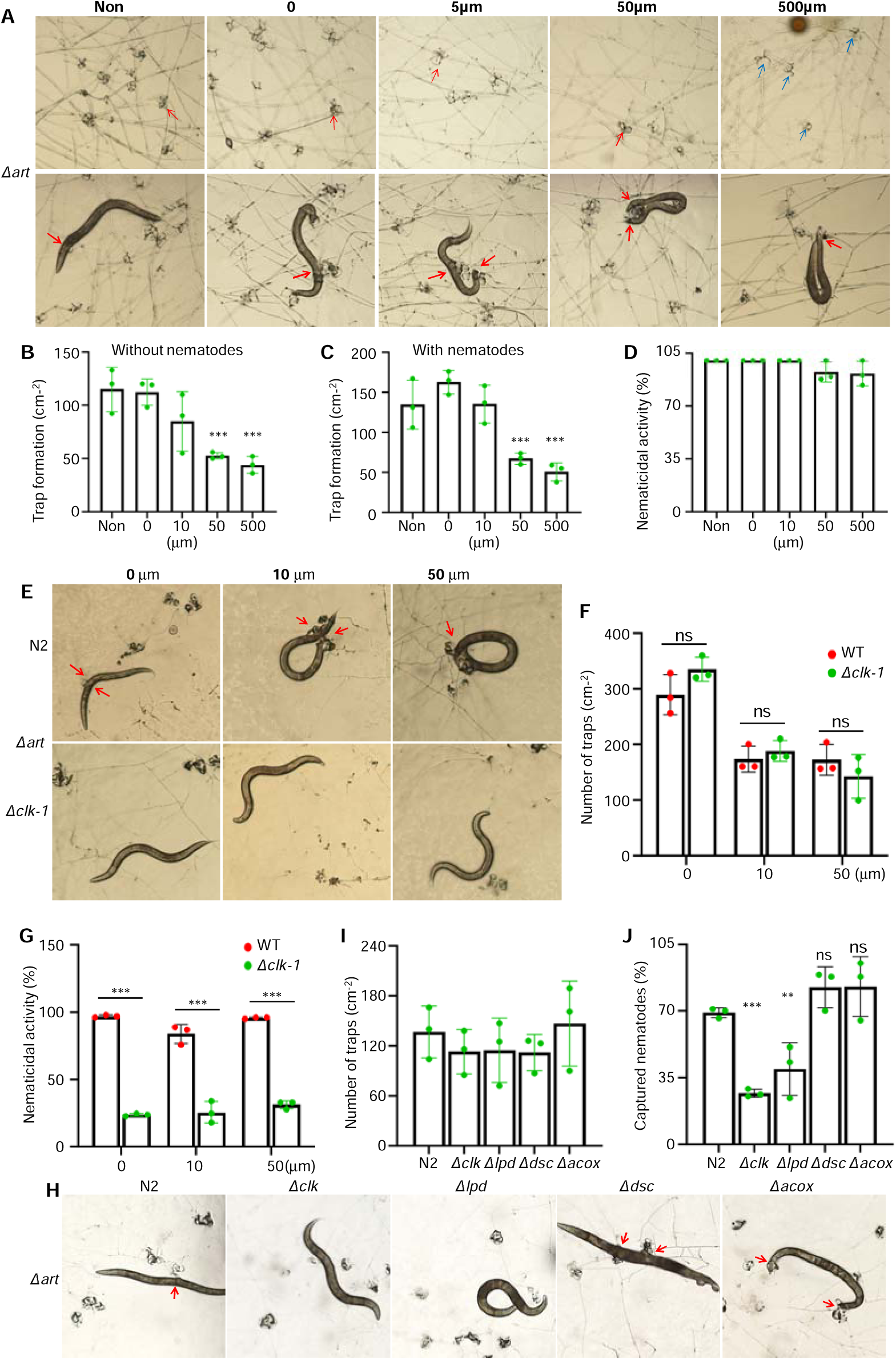
Evaluation of the effects of UQ_10_ and COQ7 on the formation of trapping devices and their nematicidal ability in the mutant. Δ*art*. (**A**) Pictures of trapping device development in the mutant Δ*art* treated without (up) and with (down) nematode treatment in the presence of different concentrations of UQ_10_. Non: control without UQ_10_ treatment. 0: with solvent only treatment. Red arrow: mature trapping devices. Blue arrow: immature trapping devices. (**B**) Comparison of trapping devices in the mutant Δ*art* treated with different concentrations of UQ_10_. (**C**−**D**) Comparison of trapping devices (**C**) and nematode-capturing ability (**D**) in the mutant Δ*art* treated with nematodes in the presence of different concentrations of UQ_10_. (**E**) Pictures of trapping device development and nematicidal ability in the mutant Δ*art* treated with two different nematodes N2 and the mutant Δ*clk-1* in presence of two different concentrations of UQ_10_. Red arrow: nematode-trapping devices. (**F**−**G**) Comparison of trapping device development (**F**) and nematicidal ability (**G**) in the mutant Δ*art* treated with two different nematodes in the presence of different concentrations of UQ_10_. (**H**) Pictures of the formation of trapping devices in the fungal mutant Δ*art* induced by five different nematodes, including nematode N2, and the nematode mutants Δ*clk-1*, Δ*lpd*, Δ*dsc*, and Δ*acox*. (**I**) Comparison of trapping devices induced by five different nematodes. (**J**) Comparison of nematicidal ability of trapping devices in the fungal mutant Δ*art* induced by five different nematodes indicated that the nematode mutant Δ*clk-1* is the last prey the trapping devices prefer to capture. Significance was tested using an unpaired t test (\**P* < 0.05, \*\**P* < 0.01, \*\*\**P* < 0.001, ns = not significant).

Interestingly, the trapping devices of the Δ*art* mutant treated with 500 μM UQ_10_ still exhibited strong nematicidal activity, although with a slight reduction compared to the other groups (Fig. 5D). These results suggest that UQ_10_ may not fully meet the fungus’s needs, even though it can inhibit trapping device development. To further evaluate the role of UQ_10_ in trapping device formation and nematicidal activity, we used the nematode mutant Δ*clk-1*, along with its WT strain N2. We found that the Δ*clk-1* mutant induced similar trapping devices in the Δ*art* fungal mutant as the N2 nematodes did (Figs. 5E–5F), confirming that the *clk-1* defect in the nematodes did not affect trapping device formation in the fungus. As expected, with or without UQ_10_, the Δ*art* mutant, like the WT strain, captured significantly more N2 nematodes than the Δ*clk-1* nematodes (Fig. 5G). However, the Δ*art* fungal mutant captured 3.1 times more N2 nematodes than Δ*clk-1* nematodes, whereas the WT fungal strain captured 2.6 times more N2 nematodes than the Δ*clk-1* nematodes, indicating that the Δ*art* fungal mutant has a much higher demand for N2 nematodes than the WT fungal strain (Figs. 5G and 3C). Furthermore, the trapping devices of the Δ*art* mutant, like those of the WT fungal strain, displayed the weakest nematode-capturing ability towards the Δ*clk-1* nematode mutant, compared to the five different nematode mutants (N2, Δ*acox*, Δ*dsc*, Δ*lpd*, and Δ*clk-1*) (Figs. 5H–5J and 3F).

### 6. Choosing *art* gene cluster over *Coq7* is associated with the origin of NTFs

We hypothesized that the loss of *coq7* and the use of arthrobotrisins in NTFs under elevated oxygen conditions might be linked to the origin and evolution of trapping devices and their predatory behavior in these fungi. To investigate this, we analyzed 21 genomes, including 9 NTFs (Orbiliomycetes), 4 fungi from its sister lineage (Pezizomycetes), 5 fungi from Eurotiomycetes (all containing the art gene cluster), 2 fungi from Saccharomycetes (which lack the art gene cluster), and 1 fungus from Taphrinomycetes (also lacking the art gene cluster) as an outgroup. Using this dataset, we constructed a time-calibrated phylogenetic tree with five calibration points using MCMCtree. Interestingly, the loss of the *coq7* gene occurred between 413 and 258 million years ago (Mya), following the acquisition of the *art* gene cluster by the ancestor of NTFs around 464 Mya (Fig. 6). This suggests that the acquisition of the art gene cluster may have reduced oxygen levels inside fungal cells, making the involvement of coq7 in respiration redundant or less essential.

**Fig 6.**
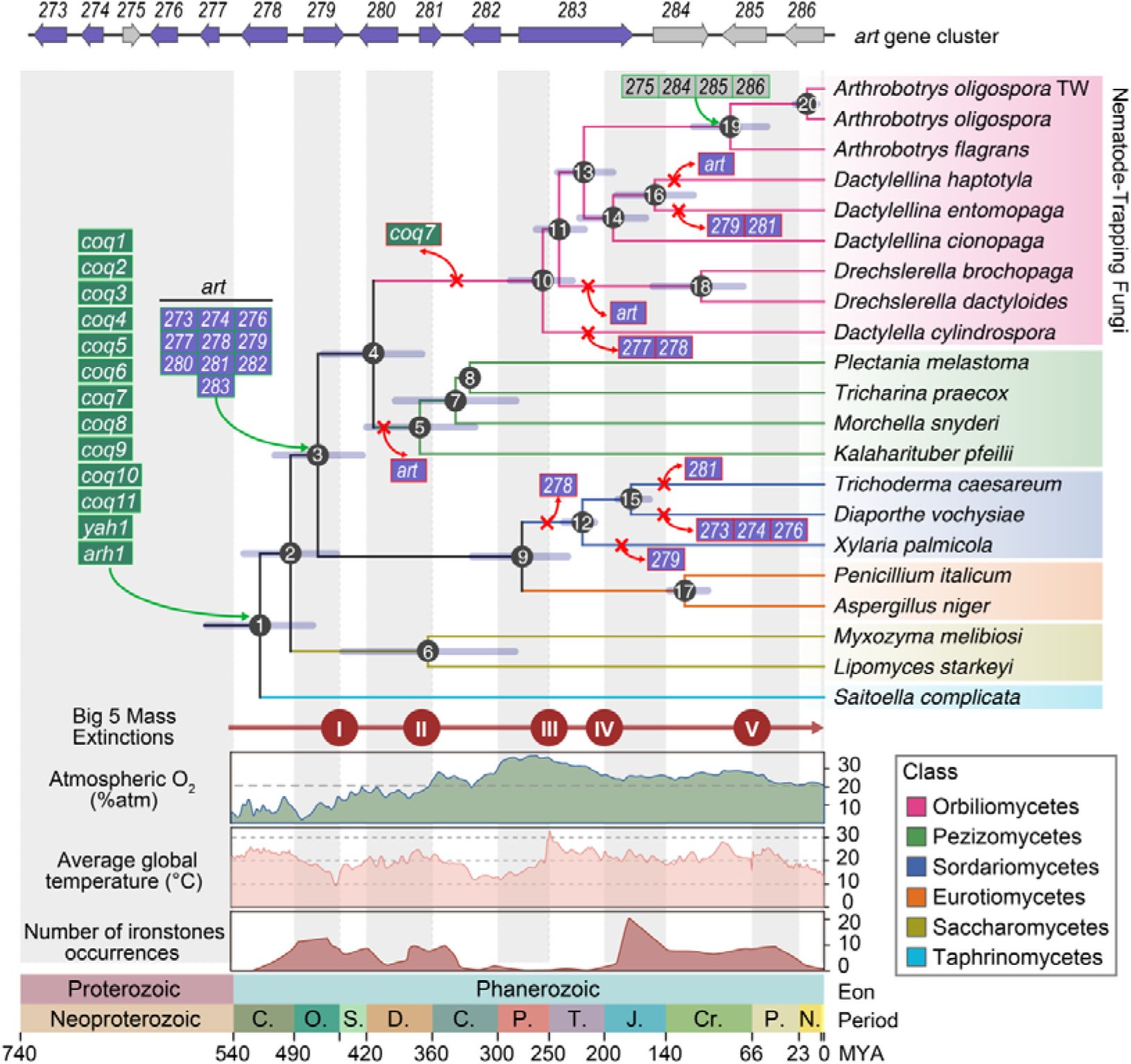
Dynamic evolutionary landscape of UQs biosynthetic genes and *art* gene cluster. Divergence times of all nodes were estimated by MCMCTree in PAML package with 278,923 4-fold degenerate sites from 1256 single copy orthologous genes and calibrated with 5 calibration points: node1 (498-571 Mya), node 4 (357-459 Mya), node 12 (216-267 Mya), node 15 (108-181 Mya) and node 17 (64-135 Mya); The error bars represent the 95% highest posterior density (HPD) of a node age; The numbered circles at each node denote the node sequence of the inner nodes; The red cross and an arrow symbol was used to indicate a certain gene loss event. The first panel represents predicted Phanerozoic atmospheric O_2_ (Data source from [8]); The second panel represents Phanerozoic global average temperature (Data source from [50]); The third panel represents the number of occurrences of Phanerozoic ironstones (Data source from [52]); The x-axis represents geological time periods spanning from the Proterozoic to the Phanerozoic Eon; The letter abbreviations denote: Cambrian (C), Ordovician (O), Silurian (S), Devonian (D), Carboniferous (C), Permian (P), Triassic (T), Jurassic (J), Cretaceous (Cr), Paleogene (P), and Neogene (N) periods.

In a previous study, we demonstrated that the formation of trapping devices in *A. oligospora* is temperature-dependent [18]. To investigate the evolutionary forces driving the acquisition of the art gene cluster and the loss of coq7 in fungi, we mapped three biogeochemical properties—atmospheric OL levels [35], average global temperature [55], and ironstone occurrences [56, 57]—onto the time-calibrated evolutionary tree. Geological data indicate that around 464 million years ago (Mya), when the NTF ancestor acquired the *art* gene cluster, atmospheric oxygen levels gradually increased from approximately 1.78% in the early Ordovician (around 477 Mya) to 19.88% in the middle Silurian (around 422 Mya) (Fig. 6). During the same period, Earth’s temperatures underwent a sharp V-shaped fluctuation, with average global temperatures rapidly dropping to 12°C and then rising to 25°C between 420 and 506 Mya. A similar, though less pronounced, trend was observed in ironstone levels during the Ordovician (Fig. 6). These findings suggest that the *art* gene cluster may have facilitated fungal adaptation to the gradually increasing oxygen levels.

Interestingly, species with fruiting bodies in the class Pezizomycetes, which share a common ancestor with NTFs in the class Orbiliomycetes, lost the *art* gene cluster rather than the *coq7* gene around 413 Mya (95% CI: 369–459 Mya) (Fig. 6). Notably, a transient decline in atmospheric oxygen levels occurred between 380 and 420 Mya. Additionally, fungal species in Orbiliomycetes and Sordariomycetes lost key genes (*Ao273*, *Ao274*, *Ao276*, *Ao277*, *Ao279*, *Ao281*) in the *art* gene cluster, or the entire *art* gene cluster, as oxygen concentrations began to gradually decline from 37.2% to present-day levels after 263 Mya (Fig. 6). This correlation between oxygen levels and the evolutionary dynamics of the *art* gene cluster suggests that environmental oxygen concentration played a crucial role in the gain and loss of genes within this *art* cluster. Further analysis of gene gains and losses related to stress responses revealed an enrichment of oxidative stress-related genes at nodes 2, 3, 4, 5, 6, and 11, corresponding to periods of rising oxygen levels (Figs. S4A–S4B and Table S2). This indicates that fungal cells likely experienced significant aerobic stress during these times.

### 7. Distribution of the *art* gene cluster in *Aspergillus* and *Penicillium*

To better understand the distribution patterns of the *art* gene cluster within the fungal kingdom, we conducted a comprehensive survey of *art* gene cluster homologs across 2,057 fungal genomes, using the *A. oligospora art* gene cluster as a search template. We retained only the hits containing *Ao283*, the core biosynthetic gene in the *art* cluster, along with at least four additional genes from the cluster for further analysis. To detect potential pseudogenized versions of the *art* gene cluster in genomes lacking a complete cluster, we performed BLAT searches. Our analysis identified analogues of the *art* gene cluster in 103 genomes, representing 86 species across 33 fungal genera. These 103 fungal genomes were distributed across several fungal classes, including Eurotiomycetes (13.2%, 52/395), Sordariomycetes (6.6%, 32/488), Dothideomycetes (5.7%, 10/174), Lecanoromycetes (27.3%, 3/11), and Orbiliomycetes (66.7%, 6/9) (Fig. S5 and Table S3).

Global fungal occurrence data from the GlobalFungi database [58] indicated that 67% of the genera (22/33) containing the *art* gene cluster are dominant fungal genera, capable of surviving across a wide range of environmental conditions and, in some habitats, thriving to become the dominant group (Table S4). Moreover, the majority of species (87%, 73/86) belonged to these dominant genera (Table S5). Further analysis of the turnover of gene members within the *art* cluster among the 86 fungal species revealed that only species in *Arthrobotrys*, along with some species in *Aspergillus* and *Penicillium*, retained an intact analogue of the *art* cluster. In contrast, other species had lost one or more key biosynthetic genes (Fig. 7). The high occurrence rates of the *art* gene cluster in dominant fungal genera, combined with the retention of intact gene clusters in *Arthrobotrys*, *Aspergillus*, and *Penicillium*, suggest that the presence of this gene cluster may play a role in the adaptation of these fungi to common environmental conditions. This is particularly relevant, as *Aspergillus* and *Penicillium* are among the most dominant fungal genera in contemporary ecosystems.

**Fig 7.**
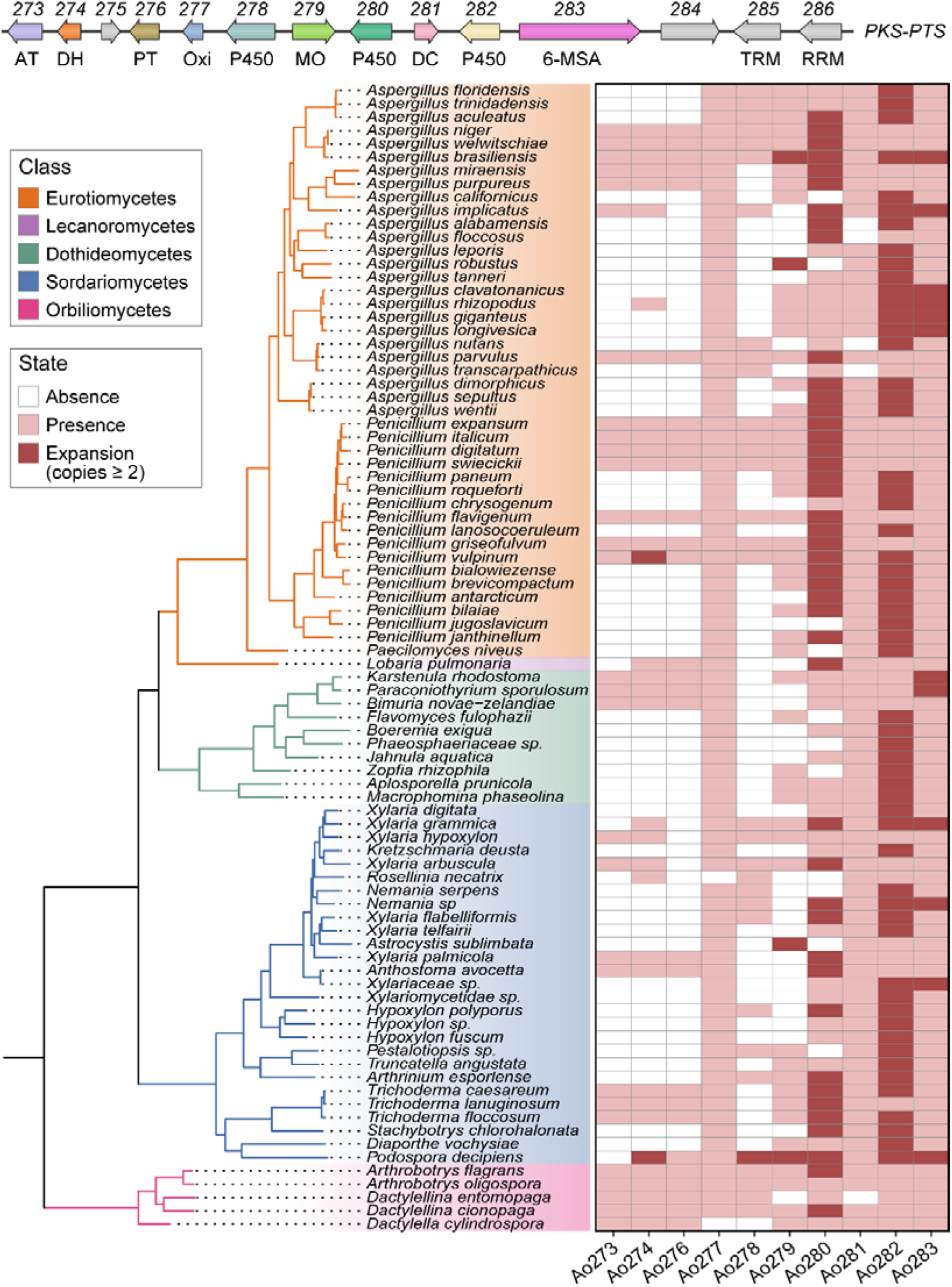
Gene members of *art* gene cluster change among various fungal species containing this gene cluster. The phylogenetic tree delineates evolutionary relationships of 86 fungal species from Eurotiomycetes (n=43), Lecanoromycetes (n=1), Dothideomycetes (n=10), Sordariomycetes (n=27) and Orbiliomycetes (n=5). These 86 fungal species were obtained by deduplicating 103 genomes containing *art* (only one genome representing the species was retained when multiple genomes of the same species contained *art*). Heatmap columns correspond to 10 biosynthetic genes (*Ao273*-*Ao274*, *Ao276*-*Ao283*) for the biosynthesis of arthrobotrinsin type of meroterpenoids within the cluster; White denotes absence, light pink denotes presence, and dark pink denotes gene expansion.

## Discussion

Numerous studies suggest that oxygen acts as a developmental morphogen in metazoan embryos, driving genomic and metabolic reprogramming during periods of oxygen fluctuation [59–62]. However, the effects of oxygen fluctuations on fungal morphogenesis, as well as on their metabolic and genomic evolution, remain largely unexplored. Previous research has shown that antioxidant enzymes and oxygen-utilizing enzymes, such as NADPH oxidases (Noxs), are upregulated by mitogen-activated protein kinases (MAPKs) under hyperoxic conditions [63, 64]. Furthermore, both NoxA and NoxB are essential for the development of fungal infection structures in certain plant pathogens [65]. However, direct evidence linking hyperoxia to the induction of fungal infection structure formation is still lacking, and the roles of specific metabolites in response to hyperoxia remain poorly understood.

We found that the formation of trapping devices and their nematicidal activities were significantly higher in the predominant NTF *A. oligospora* under normal oxygen conditions compared to low oxygen conditions. This suggests that oxygen is a key factor driving the formation of trapping devices. Surprisingly, genomic, metabolic, and functional analyses revealed that all NTFs lack the essential *coq7* gene, and *A. oligospora* utilizes UQ_8_ instead of UQ_9_-_10_ for low-aerobic respiration. Extensive genomic analysis showed that only 25 out of 2,057 fungal genomes were missing the critical coq7 gene, including all 9 NTFs from Ascomycota. Importantly, COQ7 is a membrane-bound di-iron protein belonging to a unique subset of the ferritin family [37–39], indicating its role not only in ubiquinone (UQ) biosynthesis but also in iron sequestration. Our own work has demonstrated that the trapping devices of *A. oligospora* contain a large number of iron-rich, electron-dense bodies, which is a result of the absence of the Ccc1-mediated vacuolar iron detoxification mechanism, typically conserved in most fungi. Consequently, COQ7 is chemotaxis and binds to iron-rich trapping devices due to its ferritin-like properties. Our results showed that the trapping devices preferentially captured N2 nematodes over the *coq7*-deficient nematode mutant Δ*clk-1*, suggesting that COQ7 is a key target for the NTF *A. oligospora*.

Time-calibrated evolutionary analyses suggest that the last common ancestor of NTFs within the Orbiliomycetes may have lost the *coq7* gene during the Cambrian-to-Devonian period (300–420 Mya), a time characterized by "superoligotrophy," marked by extremely low nutrient conditions [66–68]. Previous studies have proposed a link between alterations in quinone species used in respiration and lifespan. For instance, mutations in the *clk-1* gene of *Caenorhabditis elegans* lead to an extended lifespan and a general slowing of developmental rates [48, 49, 69, 70]. It is hypothesized that the chemical properties of the altered semiquinone, resulting from the absence of *coq7*, reduce reactive oxygen species (ROS)—a key determinant of aging—thereby slowing the accumulation of oxidative damage and potentially promoting longer lifespans [71]. Thus, the loss of the *coq7* gene in Orbiliomycetes may have conferred a survival advantage during the "superoligotrophy" period. Our own study further suggested that the loss and acquisition of key genes responsible for iron storage and chelation were strongly associated with temperature changes. Notably, the Ccc1-mediated vacuolar iron storage mechanism was lost during the Late Paleozoic Ice Age, while the desferriferrichrome-mediated iron storage pathway was acquired during the high temperatures of the Cambrian period [18]. It is also possible that the loss of the *coq7* gene coincided with a global temperature drop during the Late Paleozoic Ice Age (Fig. 8).

**Fig 8.**
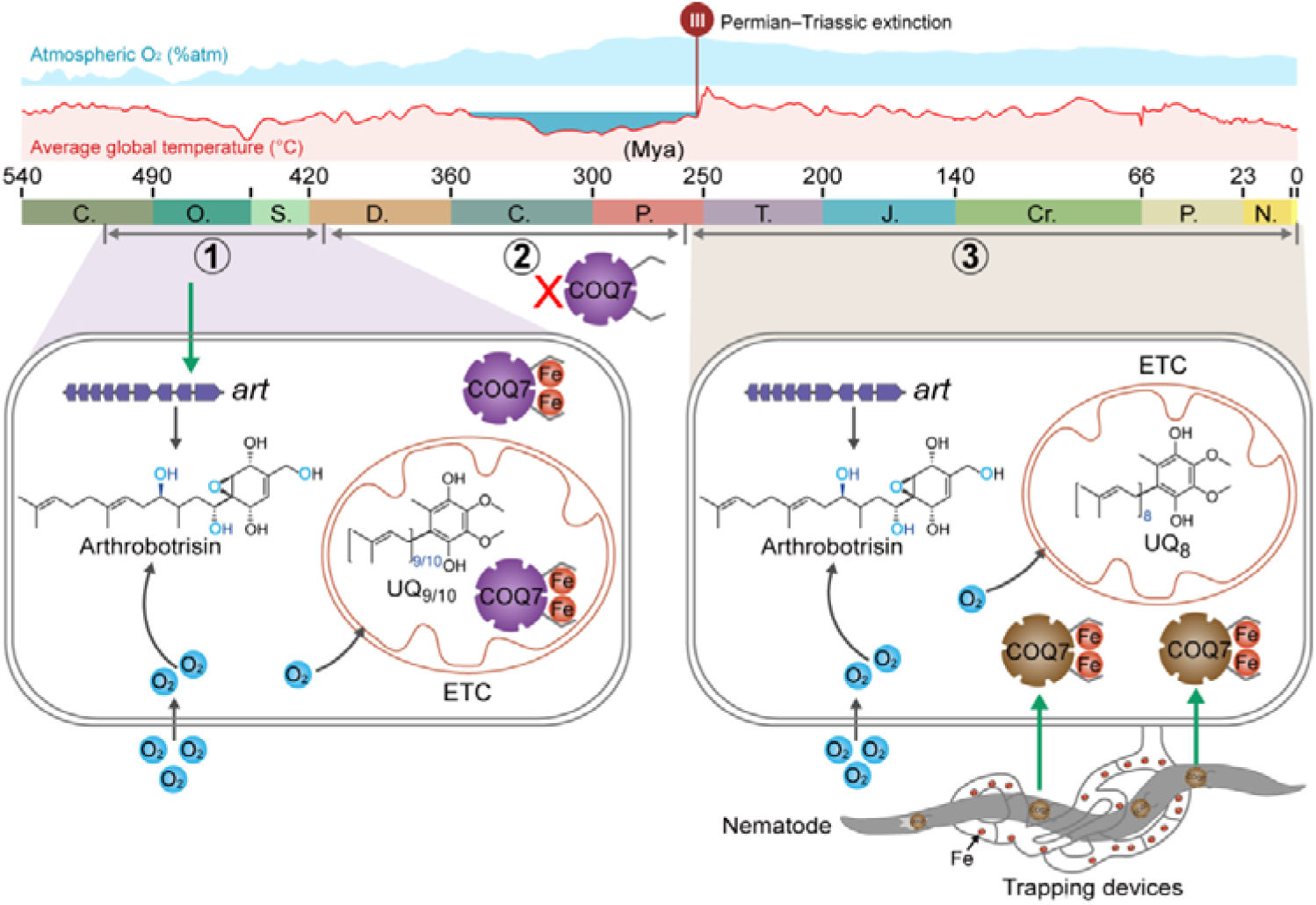
The roles of *coq7* in the origin and evolution of trapping devices and the predatory behavior of trapping devices of NTF. ETC: electron transferring chain.

The absence of the *coq7* gene in NTFs likely posed respiratory challenges as global oxygen levels increased during the Carboniferous (300 Mya) and Cretaceous (100 Mya) periods, reaching approximately 30% [72, 73]. As environmental oxygen levels rose, NTFs required COQ7 or ubiquinones (UQs) with long prenyl chains (C9–C10) to support elevated aerobic respiration and metabolic processes, as high oxygen concentrations lead to iron overload. Our results support this notion, showing that UQ_10_ strongly inhibited trapping device formation. Additionally, *A. oligospora* preferentially killed large numbers of COQ7-containing N2 nematodes, while sparing those lacking COQ7. Previous studies suggest that nematodes are typically iron-deprived animals [74, 75]. For example, *C. elegans* can detect pyoverdine, an iron siderophore secreted by *Pseudomonas aeruginosa*, and feed on the bacterial prey [76, 77]. In the predator-prey relationship between NTFs and nematodes, this dynamic is reversed. The ferritin-like properties of COQ7 in nematodes, which readily bind iron, may trigger the fungal ability to capture and prey on nematodes, due to the iron-rich nature of the trapping devices in NTFs (Fig. 8). This aligns with the dual function of the trapping devices: both attracting and capturing nematodes.

To elucidate the mechanism by which *A. oligospora* responds to high oxygen levels, we focused on the highly oxygenated arthrobotrins, which possess a triprenyl-phenol skeleton similar to UQ_3_. These compounds serve as key chemotaxonomic markers for *A. oligospora*. Triprenyl-phenols and their oxygenated derivatives are common meroterpenoids derived from phenols in fungi [78]. Notable examples include the potent antifungal yanuthones from *Aspergillus* and *Penicillium*, as well as phytotoxic macrophorins from *Macrophoma*, which causes fruit rot in apples [79]. The biosynthetic pathway of these triprenyl-phenol metabolites has only been elucidated in *A. oligospora*, due to the relatively simple metabolism and fewer biosynthetic gene clusters in *A. oligospora* [19, 20, 24–28]. Previous studies have shown that a deficiency in the arthrobotrin biosynthetic pathway significantly increases the formation of trapping devices and nematicidal activities but inhibits fungal colonization in field soils [40]. In this study, we demonstrate the functions of arthrobotrins in fungal adaptation to increased oxygen levels, as UQ_8_ in *A. oligospora* is used only under low oxygen conditions (Fig. 8). The absence of arthrobotrins greatly facilitated the accumulation of UQ_8_ and enhanced the formation of trapping devices and nematicidal activities in the Δ*art* mutant under normal oxygen conditions compared to low oxygen conditions. Notably, the Δ*art* mutant exhibited a greater demand for UQ_10_ and N2 nematodes than the WT strain. Our results in this study explain why the fungus has more trapping device, but the viability has decreased, because the trapping device is a pathological manifestation of excess iron. The limited nematicidal activity of trapping devices is also consistent with the philosophy of iron chelation treatment therapy.

Notably, the Orbiliomycetes acquired the art gene cluster around 452 Mya, prior to the loss of the key *coq7* gene around 413 Mya. Historical geological analyses reveal a gradual increase in global oxygen levels, from 1.78% in the early Ordovician (approximately 477 Mya) to 19.88% in the middle Silurian (approximately 422 Mya). Interestingly, fungal species in Pezizomycetes, which share a common ancestor with Orbiliomycetes, lost the *art* gene cluster around 413 Mya, while NTFs in Orbiliomycetes lost the *coq7* gene. This divergence in gene loss strategies suggests that the *coq7* gene and the *art* gene cluster may play similar functions in response to fluctuating aerobic conditions, owing to their structural similarities. A detailed analysis of the evolutionary history of NTFs revealed that only *Arthrobotrys* retains an intact art gene cluster, while other NTFs have either partially or completely lost this cluster. This observation aligns with the fact that *A. oligospora* is the most widely distributed NTF across diverse environments.

In conclusion, we found that oxygen plays a critical role in inducing the formation of trapping devices by increasing the availability of free iron ions. Unexpectedly, all NTFs lack the *coq7* gene, which is essential for the biosynthesis of UQs with long prenyl chains (C9–C10) and for iron chelation. The loss of the *coq7* gene is linked to the origin of NTFs during the "superoligotrophy" period. Interestingly, *A. oligospora* utilizes UQ_8_ for hypoxic respiration and a class of arthrobotrin meroterpenoids for hyperoxic respiration (Fig. 8). UQ_10_ and COQ7 strongly inhibit trapping device formation, which selectively capture COQ7-containing nematodes, suggesting that COQ7 is a key target for NTFs. Our results indicate that the loss of *coq7* played a significant role in the origin and evolution of trapping devices, and that the capture of COQ7 became an important strategy for the predatory behavior of these trapping devices (Fig. 8). Furthermore, our findings suggest that NTFs serve as an effective eukaryotic model for elucidating novel mechanisms of excess iron production and exploring iron chelation therapy.

## Data availability

Our sequences files of transcriptome are accessible from the National Center for Biotechnology Information (NCBI) under BioProject accession: PRJNA1195634 (https://www.ncbi.nlm.nih.gov/bioproject/PRJNA1195634).

