## supporting information for "Nematode-Trapping Fungus *Arthrobotrys oligospora* is Hungry for COQ7 to Alleviate Environmentally Triggered Iron Overload"

Materials and methods

**Fungal strain and culture conditions**

*A. oligospora* YMF1.3170 was obtained from the State Key Laboratory for Conservation and Utilization of Bio-Resources & Key Laboratory for Microbial Resources of the Ministry of Education. PDA (potato 200 g/L, glucose 20 g/L, agar 15 g/L), water agar (15 g/L agar) media were used for analyzing mycelial growth and related phenotypic traits. For the flask culture, strain was initially cultured on PDA medium at 28 °C for 10 days to obtain conidia, and then the conidia were inoculated in 250 mL of liquid PDB medium (potato 200 g/L, and 20 g of glucose per liter) with a final concentration of 1x10^5^ conidia/mL under shaking (180 rpm) and static conditions (0 rpm) at 28 °C.

**Acquisition and cultivation of nematode mutant strains**

The *Caenorhabditis elegans* mutant strains used in this study were obtained from the State Key Laboratory for Conservation and Utilization of Bio-Resources & Key Laboratory for Microbial Resources of the Ministry of Education. The strains, including both wild-type and mutant lines, were requested through the standard organization strain request process. Upon arrival, the nematodes were maintained at 20°C on Nematode Growth Medium (NGM) plates seeded with *Escherichia coli* OP50, as per standard cultivation protocols. The mutation status of the strains was confirmed by PCR amplification, as detailed in the strain descriptions provided by the repository. Nematode populations were synchronized as described in [80]. In brief, gravid adults were transferred onto fresh NGM plates after growing for about 2 days at 20 °C, and eggs were collected after allowing the adults to lay eggs. The eggs were then allowed to hatch overnight at 20 °C, and synchronized populations of L1 larvae were isolated. These larvae were grown to the desired developmental stage (L4) before being used for experimental purposes.

**Fungal colony growth** **under normal oxygen and low oxygen conditions**

AnnaeroPack-Anaero is a disposable oxygen-absorbing and carbon dioxide-generating agent for use in anaerobic jars. The mutant strain *∆Ao283* and the wild-type were cultured on PDA medium within anaerobic jars. Use one sachet of AnaeroPack-Anaero for a 2.5 liters anaerobic jar and discard sachet after the reaction 30min to created low oxygen conditions (approximately 10% oxygen). Smart Sensor (Model: AR8100) oxygen detector was used to detect the oxygen concentration inside the anaerobic jars. Both WT and mutant strains were cultured on PDA at 28 °C in normal oxygen and low oxygen conditions, and their growth rates and colony morphology were observed. Hyphal morphology was examined using 5-day-old PDA cultures grown at 28° C.

**Trap formation and nematicidal activity assays under normal oxygen and low oxygen conditions.**

WT and mutant strains were cultured on 9 cm agar plates under normal oxygen and low oxygen at 28 °C for 3−4 days, and then about 400 nematodes (*Caenorhabditis elegans*) were introduced to each culture. After 12 h, traps and captured nematodes per plate were observed under a microscope and counted at specific time-points. Three hundred microliters of the nematode suspension were then transferred to each Petri dish and gently mixed. All dishes were incubated at 28 °C. Nematode mortality was assessed at 12h exposure periods using a normal binocular microscope, and four visions (1cm^2^/per) in each Petri dish were picked at random for counting. Nematodes were considered to be dead when their bodies were straight, and they failed to move on physical stimuli with a fine needle. Nematicidal activity was evaluated according to the mean corrected percentage of dead nematodes. All tests were repeated three times, and the data obtained were statistically analyzed.

**Assay for iron contents.**

The WT and mutant strain were inoculated in 250 mL potato dextrose broth (PDB) flasks and cultured at 28 °C (180 rpm or 0 rpm) for 7 days. The fermentation broth was extracted with the equal volume of ethyl acetate, the mass of mycelium was weighed and recorded, then dissolved in 30 mL of distilled water and homogenized to obtain a well-ground mixture. Subsequently, 1 mL of the mixture was centrifuged at 12,000 rpm for 5 min to remove the supernatant for the following experiments. An iron assay kit (#ab83366, Abcam) was used for evaluation of the intracellular ferrous iron level (Fe^2+^ and Fe^3+^) in fungi. First, mycelia (20−80 mg) were collected in phosphate-buffered saline (PBS) and homogenized in iron assay buffer using a dounce homogenizer sitting on ice with 50−100 passes. Iron reducer was then added into the collected supernatant, thoroughly mixed, and incubated. Finally, an iron probe was added, mixed, and incubated for 1 h. the resulting solution was immediately measured on a microplate reader at OD = 593 nm.

**LC–MS/MS sample extraction and preparation**

The WT and mutant strain were inoculated in 250-ml PDB flasks cultured at 28°C under shaking(180rpm,120rpm,60rpm) and static conditions (0 rpm) for 7 days prior to analysis by high performance liquid chromatography with diode array detection MS (UPLC-DAD/MS). The fermentation broths were extracted with an equal volume of ethyl acetate, and the organic layers were evaporated and dissolved in 1 ml of menthol and filtered through a 0.22 μm filter membrane. All samples were subsequently analyzed using a HPLC system coupled with a Q Exactive Focus Orbitrap mass spectrometer (Thermo Fisher) and equipped with an Agilent Zorbax ODS 4.6 by 250 mm column (Agilent, Santa Clara, CA) in the electrospray ionization (ESI) mode. The data were acquired in both positive and negative ion modes. To ensure the accuracy of compound identification and quantification, QC samples were injected periodically throughout the analysis to monitor instrument stability and data reproducibility. The metabolites extracted from WT and mutant strains *A. oligospora* were analyzed by LC-MS following the established methods in the literature.

**LC-MS/MS data processing and analysis**

Raw MS data files generated from the LC-MS/MS system were processed using Thermo Scientific Xcalibur software for data acquisition and processing. Raw MS data were also imported into Compound Discoverer (version=3.3) software for comprehensive data analysis. Peak detection was performed using the built-in algorithm. The raw data were processed to identify and align chromatographic peaks across all samples using a reference sample. A list of features, including accurate mass, retention time, and ion intensities, was generated for each sample. Feature Finder module was used to extract relevant features from the chromatograms. The extracted features were matched to compounds using the mzCloud databases within Compound Discoverer. The software utilized both the exact mass of detected ions and their fragment ion patterns to propose possible compound identities. The ion intensities of identified compounds were normalized against total ion count (TIC). Quantitative analysis was performed to determine the relative abundance of each metabolite in the different sample groups. Statistical analysis was conducted using the software’s built-in tools, including principal component analysis (PCA) and hierarchical clustering, to visualize group separation and identify significantly different compounds across mutant Δ*art* under shaking and static conditions.

**Quantitative Real-Time PCR Analysis**

The WT and mutant strain were inoculated in 250-ml PDB flasks cultured at 28°C under shaking(180rpm) and static conditions (0 rpm) for 7 days. After induction, hyphae were collected and frozen immediately in liquid nitrogen. Total RNA was extracted from the mycelia of each sample using an RNA extraction kit (Axygen, Jiangsu, China) and then reverse transcribed into cDNA using the PrimeScriptHRT reagent kit (with genomic DNA (gDNA); TaKaRa, Kusatsu, Japan). The resulting cDNA was used as a template for RT-qPCR to analyze the mRNA expression of *Ao273-286* with specific paired primers (Table S6) and the LightCycler 480 with SYBR green I master mix (Roche, Basel, Switzerland). *β*-Tubulin was used as an internal standard. All RT-qPCR experiments were performed in triplicates. The relative transcript level (RTL) of each gene was calculated as the difference of the cycle threshold (CT) value for the gene *β*-Tubulin and *Ao273-286* under static condition and shaking condition.

**RNA-seq data analysis**

Raw reads from both WT and Δ*art* samples were examined using FastQC. To obtain clean data, fastp (v.4.10.0) [81] was used with default parameters to filter adaptor sequences and remove low-quality reads. The reference genome and gene model annotation files for A. oligospora ATCC 24927 were downloaded from NCBI’s Genome Browser. Genome indexing was performed using ‘hisat2-build’ in Hisat2 (v.2.2.1) [82], incorporating splice sites and exon information. The filtered paired-end reads were aligned to the reference genome using Hisat2. Gene abundance for each sample was estimated using featureCounts (v.2.0.3) [83] from the aligned BAM files. Differential expression analysis was conducted using the DESeq2 R package (v.1.30.1) [83]. DESeq2 employs statistical tests to identify differential expression in digital gene expression data based on the negative binomial distribution. Resulting p-values were adjusted using the Benjamini–Hochberg approach to control the false discovery rate.

**Construction of *coq7* heterologous expression strain (E-*coq7*) using *A. oligospora***

To construct a heterologous expression plasmid, the PUC19-1300-D-HYB (PUC19) plasmid was amplified using fusion polymerase chain reaction (PCR) with Phanta Max SuperFidelity DNA Polymerase (Vazyme, Nanjing, China), following the manufacturer's instructions. The primers used in this study are listed in Table S7. The heterologous fragment was amplified from *Saccharomyces cerevisiae* BJ5464 and purified. The *coq7* gene was inserted into the PUC19 plasmid, which contains the strong *PtrpC* promoter and *TtrpC* terminator from *Aspergillus niger*. The *coq7-*plasmid and blank plasmid transformed into *A. oligospora* protoplasts, respectively. After 2−4 days of incubation at 28°C, transformation colonies were selected and transferred to new PDA plates. Genomic DNA from putative transformants was extracted after 5 days of incubation at 28°C, and PCR was performed to confirm the integration of the target gene into the genome. The mutant was identified and verified by PCR and sequencing analysis. The WT strain and E-*coq7* mutant were cultured on PDA medium at 28°C for 6 days. Vegetative hyphae were harvested from 9-cm Petri dishes, which were then incubated at 28°C for 3 and 5 days. Following induction with nematode, the hyphae were collected and immediately frozen in liquid nitrogen. Total RNA was extracted from all samples using the AxyPrep Multisource RNA Miniprep Kit (Axygen, Jiangsu, China). This extracted RNA was reverse transcribed into cDNA using the FastQuant RT Kit with gDNase (Takara, Kusatsu, Japan). The cDNA was used as a template to analyze the expression of the *coq7* gene, with cDNA from *S. cerevisiae* BJ5464 serving as the positive control template.

**Phylogenetic tree construction of 2057 fungal genomes using BUSCO**

To construct a phylogenetic tree for 2057 high-quality fungal genomes downloaded from the JGI MycoCosm database (Access: 2023-6-15) [84], we employed BUSCO (v.3.0.2) [85] to identify the Universal Single-Copy Orthologs across the genomes using the fungi_odb10 lineage dataset in BUSCO database. The identified orthologous sequences from each genome were extracted from the respective BUSCO output directories. For each ortholog group, multiple sequence alignments were performed using MAFFT (v.7.505) [86]. These alignments were rigorously inspected for quality and subsequently concatenated into a comprehensive alignment matrix. This concatenated dataset provided a holistic view of the conserved genetic information across the sampled genomes. Maximum likelihood (ML) tree with 1000 bootstrap iterations were built using FastTree (v. 2.1.11) [87].

**Distribution pattern of UQs biosynthesis genes in the fungal kingdom**

At least 13 proteins (COQ1–COQ11, YAH1, and ARH1) are necessary for the synthesis of UQs. To elucidate the distribution patterns of these proteins in the fungal kingdom, protein sequences of COQ1–COQ11, YAH1, and ARH1 from *Saccharomyces cerevisiae* were used as queries. These sequences were used to search for orthologs via BLASTp (v.2.11.0+) [88] against a custom database containing all predicted protein sequences from 2057 high-quality fungal genomes downloaded from the JGI MycoCosm database. Our criteria for BLASTp were bit scores greater than 50, minimum alignment length of 50 amino acids and minimum sequence similarity of 30%. Presence and absence of each gene were mapped to the phylogenetic tree (containing 2057 fungal genomes) to visualize the distribution of UQs biosynthesis genes in fungal kingdom.

**Identification homologue of *art* gene cluster in the fungi kingdom**

Biosynthetic gene clusters for each genome were annotated by antiSMASH 7.0 (v.7.0.0) [89] via command line version with default parameters, except for the argument ‘--taxon fungi’. In total, 84,476 secondary metabolism gene clusters were annotated from 2057 fungal genomes. A local database was generated based on all the gene clusters using the ‘cblaster makedb’ module (v.1.3.18) [90]. To identify *art* gene cluster homologous in fungal kingdom, the search function of cblaster was used to query amino acid sequences of 14 genes in *art* gene cluster (*Ao273*–*Ao286*) against the local database constructed above. Only resultant hits containing Ao283, the core biosynthetic gene of *art*, as well as at least four other genes in this cluster, were considered as a putative *art* homologues. We confirmed 103 fungal genomes containing *art* gene clusters homologues. The distribution pattern of presence and absence of *art* gene cluster was mapped to the phylogenetic tree (containing 2057 fungal genomes) to visualize the distribution of the gene cluster in the fungi kingdom.

**Gene members of *art* gene cluster among various fungal species**

The 103 fungal genomes containing the *art* gene cluster were used for this analysis. When a species had multiple genomes containing *PKS-TPS* gene cluster, only the genome with the highest number of gene members in this gene cluster was retained as the representative of that species. We identified 86 fungal species containing the *PKS-TPS* gene cluster. Evolutionary relationships of these 86 fungal species were extracted from the phylogenetic tree, which included 2057 fungal genomes. A heatmap was generated to illustrate the presence, absence, or expansion (gene copies > 2) of these genes across these fungal species.

**Phylogenetic tree construction for nematode-trapping fungi in Ascomycota**

To build a robust phylogenetic framework for exploring the evolutionary origin of trapping structures in nematode-trapping fungi, we selected 21 genomes including 9 nematode-trapping fungi (Orbiliomycetes), 4 fungi from its sister lineage (Pezizomycetes), 5 fungi (all of these contain *art* gene cluster) evenly sampled across Eurotiomycetes, 2 fungi (without *art* gene cluster) from Saccharomycetes as well as 1 fungus (without *art* gene cluster) from Taphrinomycetes as an outgroup. All-versus-all BLASTP results of 207,036 protein sequences from the 21 fungal genomes were input into OrthoFinder (v.2.5.2) [91] for gene clustering, in which the MCL algorithm [92] was enabled by setting the inflation factor to 1.5. In total, 1256 sets of 1:1 orthologous protein coding genes were identified among the 21 fungi species. Amino acid sequences of each orthogroups were aligned with with the ‘auto’ parameter, and PAL2NAL (v14) [93] was used under default parameters to create a codon alignment from MAFFT-aligned amino acids. The aligned orthogroups amino acid sequences and its codon alignment were concatenated and used to construct an amino acid (AA) and a nucleotide (NT) data matrix, respectively. Based on AA data matrix, we determined the best model using the “-m TEST” parameter, which automatically estimates the best fitting model of substitutions according to Bayesian Information Criterion values (BIC) using IQ-TREE2 (v. 2.2.2.7) [94]. A maximum likelihood (ML) tree with 1000 bootstrap iterations was built based on the best fitting model.

**Function annotations of homologous groups**

To assign function annotations to each homologous group (OG), we selected a representative sequence for each group. First, we ordered the sequences within each OG by their lengths using the ‘sortbylength’ option in USEARCH11 (v.11.0.667) [95]. Next, we employed the UCLUST algorithm to cluster the sequences within each OG based on 30% amino acid sequence similarity. The sequence with the highest abundance in each OG was designated as the representative sequence. In cases of equal abundance, the longest sequence was chosen as the representative. For each representative sequence, gene function and gene ontology (GO) were predicted using the PANNZER2 web server [96].

**Estimating divergence times of nematode-trapping fungi in Ascomycota**

To estimate the divergence times, fourfold degenerate (4D) sites were extracted using get4foldSites (<https://github.com/brunonevado/get4foldSites>) and concatenated into a single supermatrix using ‘concat’ in seqkit (v2.4.0) [97]. The Bayesian relaxed molecular clock approach was employed to estimate species divergence time using MCMCTREE in PAML (v.4.10.0) [98] (clock = independent rates, burn-in = 10,000, sample number = 100,000, sample frequency = 10, model = JC69) based on 278,923 4-fold degenerate sites from 1256 single copy orthologous genes and calibrated with 5 calibration points: node1 (498-571 Mya) [99], node 4 (357-459 Mya) [100], node 12 (216-267 Mya) [101], node 15 (108-181 Mya) [101] and node 17 (64-135 Mya) [101]. We used Tracer (v.1.7.1) [102] to visually check convergence of parameters across replicate MCMC chains in report of the MCMC runs. All the resulting effective sample sizes (ESS) ranged from 291 to 1306, suggesting that the parameters across replicate MCMC chains reached convergence (The effective sample size (ESS) should be higher than 200).

**Dynamic landscape of gene members in *art* gene cluster during the evolutionary timeline**

Based on the time-calibrated phylogenetic tree, we mapped three representative biogeochemical properties—atmospheric O_2_ [35], average global temperature [55], and the number of ironstone occurrences [56, 57] during the Phanerozoic eon onto the tree. Gene members of *art* gene cluster (*Ao273*–*Ao286*) and UQs synthesis-related genes of 21 genomes in phylogenetic tree were also mapped onto the tree to display their loss and gain at different nodes.

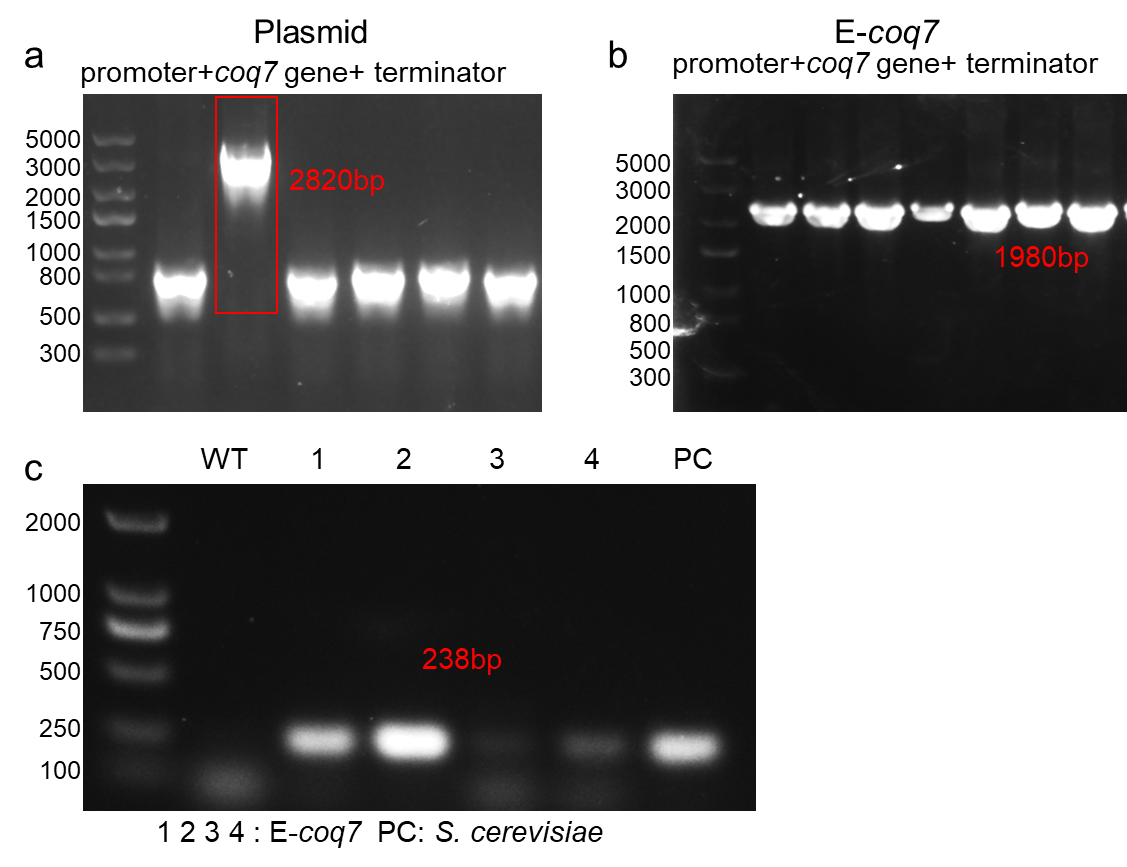
**Figure S1. Construction and verification of *A. oligospora* mutant strain E-*coq7* containing an exogenous gene *coq7* from yeast.** (**a**) Gel electrophoresis image of PCR verification of *coq7*-containing plasmid. (**b**) Gel electrophoresis image of PCR verification of *coq7*-containing transformants. (**c**) Transcriptional levels of the mutant E-*coq7.* The bands in the gel electrophoresis image represent the cDNA of the mRNA reverse transcription of the *coq7* gene in the mutant E-*coq7*.

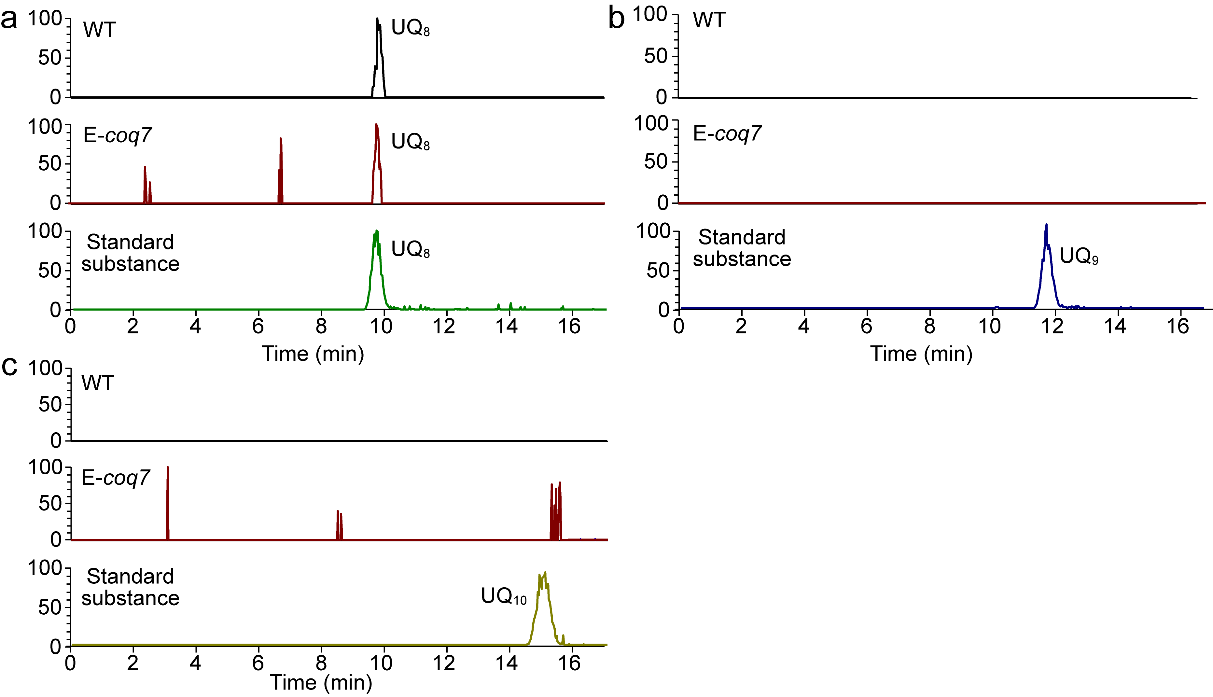
**Figure S2. Metabolic analysis of UQ_8_** (**a**)**, UQ_9_** (**b**)**, and UQ_10_** (**c**) **in the mutant E-*coq7* and WT.**

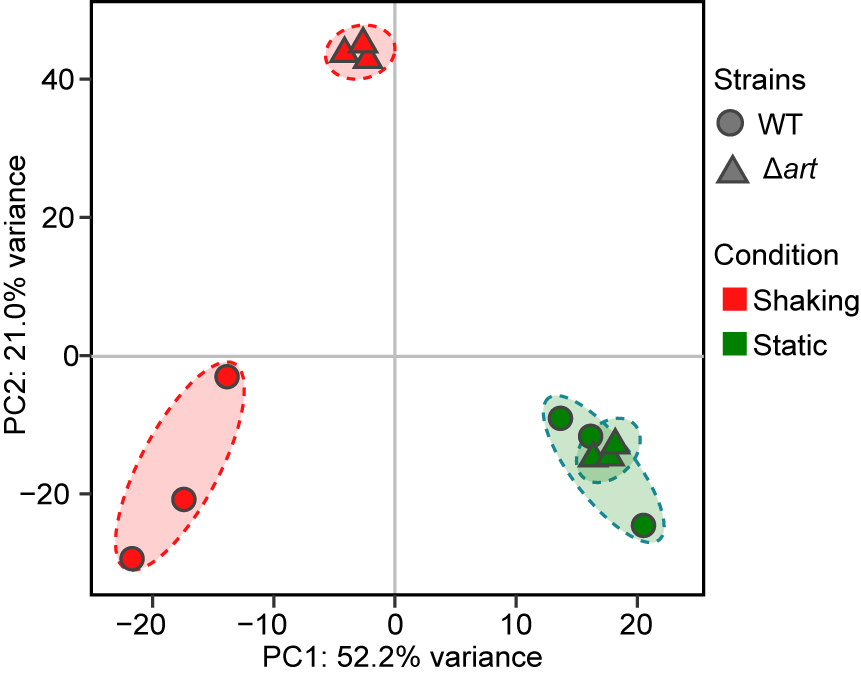

**Figure S3.** Principal component analysis (PCA) of the metabolic profiles of WT and Δart mutant strains under shaking and static condition.

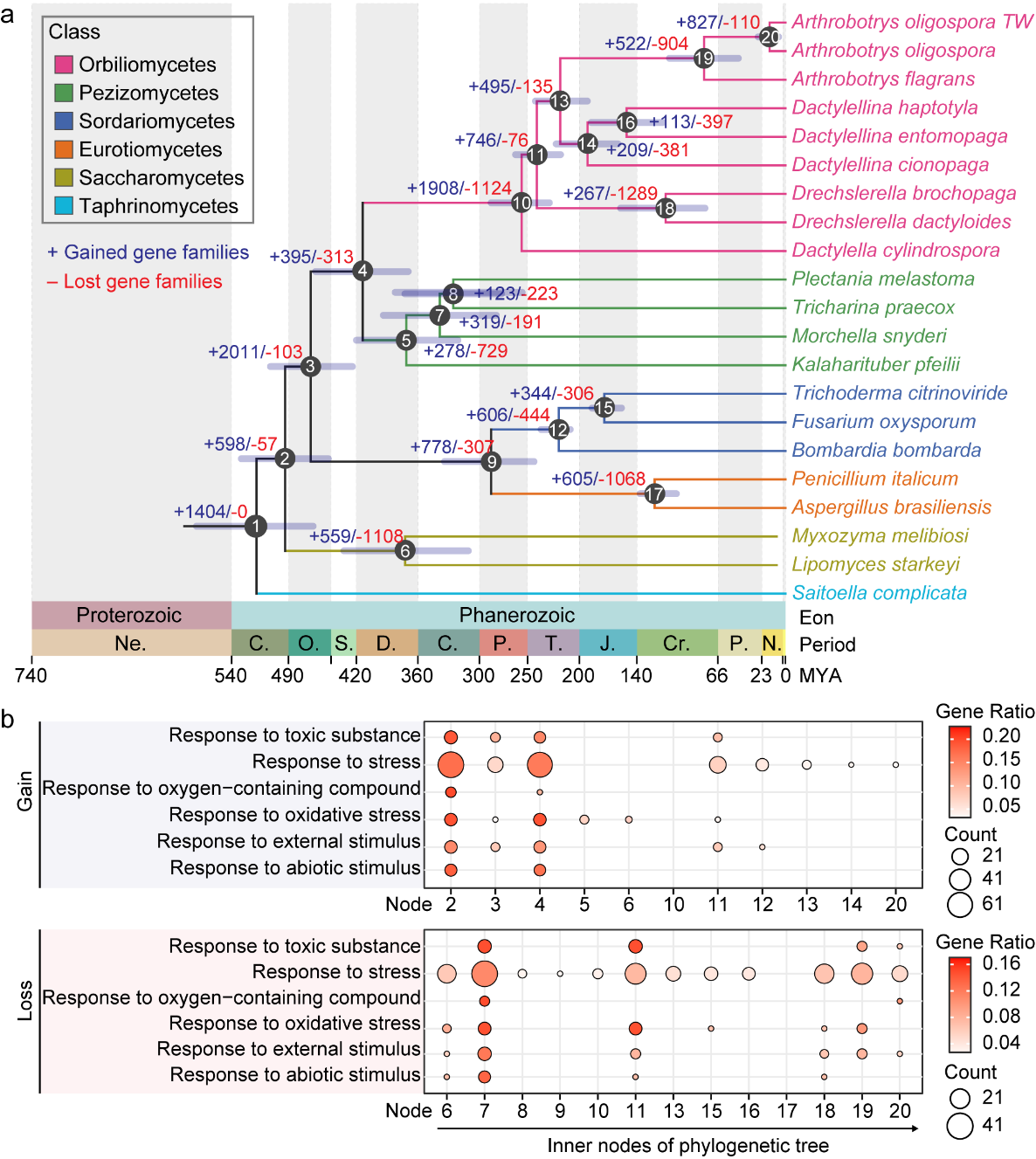

**Figure S4.** Reconstruction of ancestral gene families gains and losses during fungal evolution. (**a**) Gains and loss of all gene families in inner nodes of phylogenetic tree. Ancestral gene family sizes were estimated using the COUNT program with Dollo parsimony. Gain and loss events were indicated at each lineage of the tree, with "+" symbols representing gains and "−" symbols indicating losses. (**b**) Gains and loss of genes involving in response to stress. The node number corresponds to the node number on the evolutionary tree.

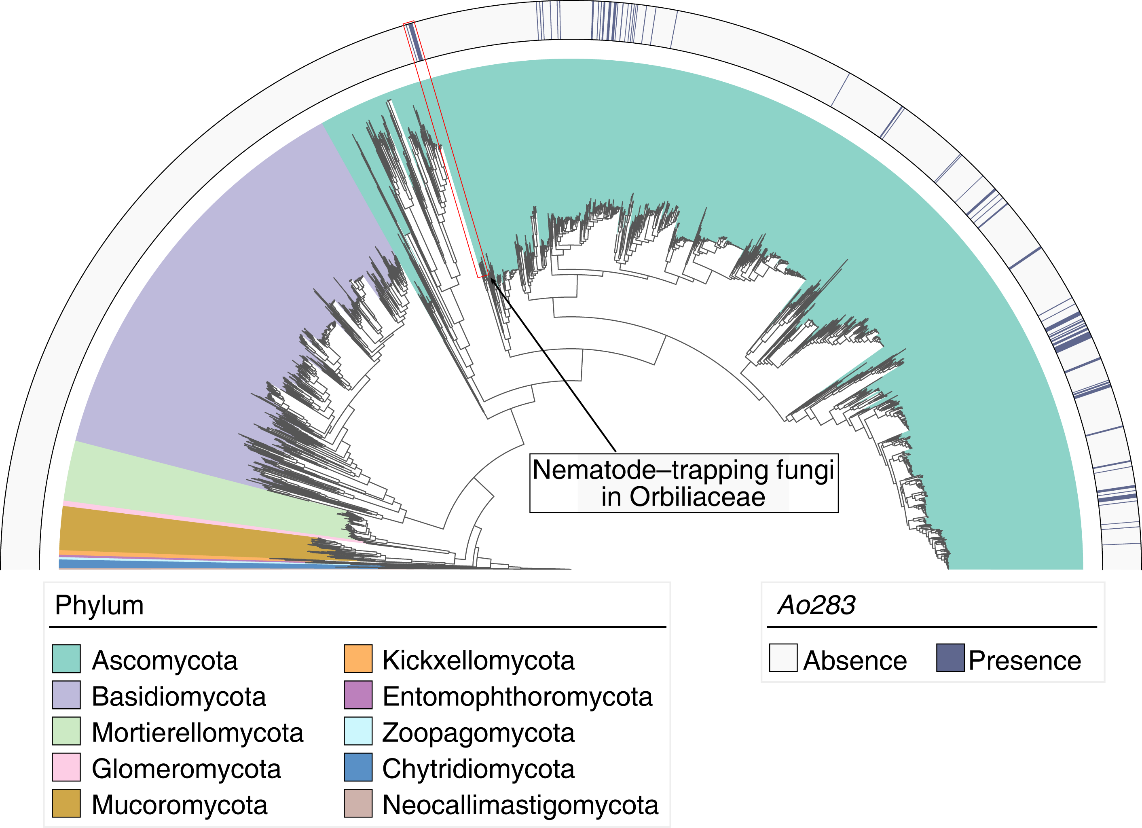

**Figure S5.** Distribution and variation of *art* gene cluster across the fungal kingdom. Presence and absence of *art* gene cluster in different fungal species across different phyla. The phylogenetic tree was constructed using 750 universal single-copy marker genes extracted from 2057 high-quality fungal genomes downloaded from the JGI MycoCosm database. The fungal groups analyzed including Ascomycota (1361 genomes), Basidiomycota (540 genomes), Mucoromycota (57 genomes), Glomeromycota (7 genomes), Mortierellomycota (67 genomes), Kickxellomycota (6 genomes), Chytridiomycota (11 genomes), Neocallimastigomycota (2 genomes), Entomophthoromycota (4 genomes), and Zoopagomycota (2 genomes). Highlighted section using red box denotes nematode-trapping fungi within Orbiliaceae. Circle outside the tree represents the presence and absence of *art*.

**Table S6**. **The primers were used for qPCR**

| **Gene** | **Primer name** | **Sequence（5' to 3'）** |
| --- | --- | --- |
| *β-tubulin* | *β-tubulin F* | GCCACCTTCGTCGGTAACTC |
|  | *β-tubulin R* | TCGTCCATACCCTCACCAG |
| *Ao273* | *qP273-F* | CGCCTGGAGTATTCGGAATGTC |
|  | *qP273-R* | GCTACCGAAGATTAGGGAAGAGAC |
| *Ao274* | *qP274-F* | TCGTACCCCCTCTGAAGGAG |
|  | *qP274-R* | TACTTCCCGTTACCCTCGGT |
| *Ao275* | *qP275-F* | GGTTGTCTGCCACCCAAAAC |
|  | *qP275-R* | TCGGCTGCTTAACATTCCGA |
| *Ao276* | *qP276-F* | TTGTTGTCGTTTACTGCGCG |
|  | *qP276-R* | CAACCCCCATACACAGACCC |
| *Ao277* | *qP277-F* | CATACCTGTCGCCTCCTTGG |
|  | *qP277-R* | AACGGCTGATACCCAAGCAA |
| *Ao278* | *qP278-F* | TACAAAGACCAACGCGTCGA |
|  | *qP278-R* | ACCAACGGGATGGCTTGAAT |
| *Ao279* | *qP279-F* | ATGCCTCATTGGAGTGGGTG |
|  | *qP279-R* | GTCCGAGTATATTGCCGCCA |
| *Ao280* | *qP280-F* | CAAGCCAAATTTCCCCACGG |
|  | *qP280-R* | GCTTCGACGGTACCTGGAAA |
| *Ao281* | *qP281-F* | GTTCACCACCACTTTGTGCC |
|  | *qP281-R* | TCCATCAGAGTCTCGCTGGA |
| *Ao282* | *qP282-F* | CAAACTTGGAACGGGCACTG |
|  | *qP282-R* | CATATGATGGCGGTCGGTCA |
| *Ao283* | *qP283-F* | TGGAATGGGTTGGGTATGGC |
|  | *qP283-R* | AGGCCCTGAAAGCTTCATCC |
| *Ao284* | *qP284-F* | GATTTCCTAGAGGCGACGGG |
|  | *qP284-R* | ATACTCCAACTGCCAGGTGC |
| *Ao285* | *qP285-F* | TCCCTTCAACTTCGACACCG |
|  | *qP285-R* | GGGACAGAAGGTGGGTTGAG |
| *Ao286* | *qP286-F* | GATCATGCTGCCCGGTATGA |
|  | *qP286-R* | GCATTCCAGGTCTGAGCTGT |

**Table S7. The primers were used for construction of mutant E-*coq7***

| **Gene** | **Primer name** | **Sequence（5' to 3'）** |
| --- | --- | --- |
|  | *coq7*-cDNA-F | CCTATTCTACCCAAGCATCGATATGTTTCCTTATTTTTACAGACGAG |
|  | *coq7*-cDNA-R | CACGAGCTACTACAGATCCCCGTTAAATTCTTTCGGCACTCCATATAG |
|  | *coq7*-plasmid-F | CTATATGGAGTGCCGAAAGAATTTAACGGGGATCTGTAGTAGCTCGTG |
|  | *coq7*-plasmid-R | CTCCTTCAATATCATCTTCTGTCGAATCCGTTTAAACGGCGCG |
|  | Ptrpc-F | CGCGCCGTTTAAACGGATTCGACAGAAGATGATATTGAAGGAG |
|  | Ptrpc-R | CTCGTCTGTAAAAATAAGGAAACATATCGATGCTTGGGTAGAATAGG |
|  | *coq7*-yz-F | TCCCTTCAACTTCGACACCG |
|  | *coq7*-yz-R | GGGACAGAAGGTGGGTTGAG |
|  | *coq7*-RT-F | ACGTCGTTATCTTCTCCTCCA |
|  | *coq7*-RT-R | ACACGGTCCAAAAATGCAGC |
